# The Regulation of Developmental Diapause is Coordinated with Molting and Olfaction

**DOI:** 10.1101/2020.03.30.016311

**Authors:** Heather R. Carstensen, Reinard M. Villalon, Johnny Vertiz, Navonil Banerjee, Elissa A. Hallem, Ray L. Hong

**Affiliations:** California State University, Northridge Department of Biology 18111 Nordhoff Street Northridge CA, 91330-8303, USA; University of California, Los Angeles Dept. of Microbiology, Immunology & Molecular Genetics and Molecular Biology Institute 615 Charles E. Young Drive S. Los Angeles, CA 90095, USA

**Author notes:** Author contributions: HRC, RMV, NB, and RLH designed and performed the experiments; JV performed the experiments; HRC, RMV, JV, NB, and RLH analyzed data; HRC, RV, and RLH wrote the manuscript.

**Keywords:** dauer larva, evolution of development, nematodes, steroid hormones, cuticle exsheathment, olfaction

## Abstract

Developmental and behavioral plasticity allow animals to prioritize alternative genetic programs during fluctuating environments. Behavioral remodeling may be acute in animals that interact with host organisms, since reproductive adults and the developmentally arrested larvae often have different ethological needs for chemical stimuli. To understand the genes that coordinate development and behavior, we used the nematode model *Pristionchus pacificus* to characterize mutants that inappropriately enter developmental diapause to become dauer larvae (Daf-c). We found several key olfactory differences between *P. pacificus* and *C. elegans* Daf-c dauers. In addition, the two *P. pacificus* Daf-c alleles disrupt steroid synthesis required for proper regulation of the conserved canonical steroid hormone receptor DAF-12, whose dauer-constitutive and cuticle exsheathment phenotypes can be rescued by the feeding of Δ7-dafachronic acid. One allele, *csu60*, has a deletion in the sole HydroxySteroid Dehydrogenase (HSD) in *P. pacificus.* Both *hsd-2(csu60)* adults and dauers show enhanced attraction to a beetle pheromone, possibly due to the heterochronic activation of dauer-specific neuronal development in the adults. Surprisingly, this enhanced odor attraction acts independently of *daf-12*, revealing unexpected targets of steroid hormones regulating ecdysis and olfaction in *P. pacificus*.

**Author Summary:** The remarkable evolutionary success of nematodes can be attributed to their dispersal ability as stress-resistant dauer larvae and the equivalent parasitic infective larvae. The decision to enter dauer development is regulated by a conserved steroid hormone receptor that receives multiple external and internal cues, though the extent in which these cues also coordinate other physiological and behavioral processes is not well understood in divergent species. We used the insect-associated nematode *Pristionchus pacificus* to genetically dissect two mutants that form dauers inappropriately, and found that one mutation in a steroidogenic enzyme, *Ppa-hsd-2*, is predicted to abolish the biosynthesis of the hormones needed to negatively regulate dauer entry when food is available. Unexpectedly, *Ppa*-HSD-2 is also required to properly complete the dauer larval molt, known as exsheathment, as well as to confer differences in adult versus dauer larvae chemotaxis behavior towards a host pheromone. Given that dauers are the host-seeking stage of parasitic nematodes, hormonal disruption represents a tenable target for biological control.

## Introduction

Organisms integrate multiple sensory stimuli continuously from their environment to make key developmental decisions and engage in context-dependent behavior. When environmental conditions are unfavorable, a portion of the population can stop feeding and suspend development. In nematodes, of the several larval stages that can undergo developmental arrest [1, 2], the dauer larvae (DL) in *Caenorhabditis elegans*, as well as the equivalent infective juvenile (IJ) or infective third-larval (iL3) stages in parasitic nematodes, undergo spectacular physiological, morphological and neuroanatomical changes [3–5]. At the gene expression level, a number of the dauer entry changes are reversible while some become permanent life history traits that mark dauer passage [6, 7]. Entry into the dauer stage is primarily determined by environmental factors, particularly the depletion of food, elevated temperature, and high levels of ascaroside dauer pheromones in response to overcrowding (Cassada and Russel, 1975; Golden and Riddle 1982 and 1984a; Jeong et al., 2005; Butcher et al., 2007). The DL is thus a striking example of developmental plasticity that is becoming more amenable to genetic analysis in diverse species of nematodes as a result of comparative studies in non-*Caenorhabditis* species [8–10].

The commitment to dauer entry requires external as well as internal factors. Multiple parallel-acting signaling factors in the dauer regulation pathway have been characterized in *C. elegans*, such as the recognition of pheromones by G-protein-coupled receptors in the chemosensory amphid neurons and signal transduction to the DAF-11/guanylyl cyclase pathway [11]. Favorable conditions also lead to signaling in the DAF-7/TGF-ß and DAF-2/insulin-like pathways [11–14]. This signaling facilitates the production of Δ4- and Δ7-dafachronic acid from the cholesterol precursor, which act as ligands that bind to the highly conserved nuclear hormone receptor DAF-12 and prevent DAF-12 from repressing genes necessary for development to reproductive adulthood [11,15–17]. Less favorable conditions produce pheromones that reduce signaling in these pathways, subsequently shifting DAF-12 from its inactive ligand-bound form to its active, ligand-free form that specifies dauer development. Despite being a non-feeding and stress-resistant stage, the DL are actively motile and receptive to external chemosensory and mechanical cues. For both parasitic and free-living nematodes, DL share the ability to survive harsh conditions but remain receptive to semiochemicals from their environment.

The commitment to dauer also prompts the rewiring of the sensory nervous system in *C. elegans* [6,18,19], but the molecular basis of this process remains poorly understood [20]. Comparisons of chemosensory behaviors in reproductive free-living adults versus developmentally arrested DL or iL3s have revealed that responses to some chemosensory cues change dramatically across life stages. For example, *C. elegans* well-fed adults are repelled by carbon dioxide (CO_2_), while *C. elegans* DL are attracted to CO_2_ [21, 22]. However, most genetic studies of chemosensory behavior have been conducted on young adults in free-living nematodes [23–25], while chemotaxis behaviors in parasitic nematodes were profiled primarily with IJs or iL3s [26–31].

To better understand the genetic network that coordinates dauer entry with development in species that evolved in fluctuating environments with hosts, the entomophilic nematode *Pristionchus pacificus* has been cultivated as a genetically tractable comparison to *C. elegans*. In the wild, *P. pacificus* has been found as DL on various species of beetles, which upon the hosts’ death, exit the dauer state and resume reproductive development to feed on the bacteria [32–34]. Thus, *Pristionchus* DL are considered to be the host-seeking stage, with adaptations that include an oily coat that promotes aggregation, survival up to a year (perhaps due to the host life cycle), and mouth form plasticity, in which dauer passage increases the proportion of the bacteriovorous stenostomatous form with faster pharyngeal pumping [35–38]. Furthermore, adult *Pristionchus* species have distinctive chemosensory preferences for high molecular weight insect pheromones and plant volatiles that differ significantly from *Caenorhabditis* species, and also differ across *Pristionchus* species [39]. However, due to the low number of DL that can be induced by starvation in the reference strain, the chemosensory profiles of *P. pacificus* DL have not yet been determined.

In *P. pacificus,* several dauer formation defective (Daf-d) mutants that fail to form DL under inducing conditions have been isolated by unbiased forward genetics as well as reverse genetics. These studies have identified genes encoding for the nuclear hormone receptor DAF-12 and the forkhead transcription factor DAF-16 as important regulators of dauer formation [8, 9]. However, while the DAF-12 and DAF-16 functions are conserved between *C. elegans* and *P. pacificus*, mutations in other Daf-d orthologs suggest divergence in the factors that act upstream of these proteins. In *C. elegans*, high population density induces the secretion of dauer pheromones consisting of a modular library of ascarosides [40–43], and mutations in the ascaroside biosynthesis enzyme, DAF-22, result in the Daf-d phenotype [44]. Interestingly, the *daf-22* gene underwent a recent duplication in *P. pacificus* that resulted in subfunctionalization in ascaroside production, yet neither mutations in the *Ppa-daf-22.1*/*daf-22.2* paralogs alone or in combination resulted in defects in dauer formation [45]. These studies indicate a possible significant divergence in *P. pacificus* and *C. elegans* dauer regulation upstream of the conserved DAF-12 and DAF-16 modules.

In contrast to the Daf-d mutants, no Dauer formation constitutive (Daf-c) mutants have been characterized extensively in *P. pacificus*, despite numerous Daf-c orthologs in the genome. Putative homologs of the *C. elegans* DAF-11/guanylyl cyclase, DAF-7/TGF-ß ligand, and DAF-2/insulin receptor have been identified in the *P. pacificus* genome (www.wormbase.org), but it remains unknown if any of these homologs are involved in dauer regulation. At the same time, targeted mutations in *P. pacificus* orthologs of other *C. elegans* Daf-c mutants did not result in dauer formation phenotypes. Specifically, mutations in the *P. pacificus* heat shock protein *Ppa-*DAF-21*/*HSP90 and *Ppa-*DAF-19/RFX transcription factor affected ciliogenesis but did not result in a Daf phenotype [46, 47]. Copy number variations in the *P. pacificus dauerless* locus, a fast-evolving gene with homologs only in the *Pristionchus* genus, control responsiveness to dauer pheromones but do not cause a Daf-c phenotype [48]. Finally, because cholesterol is a precursor for the synthesis of steroid hormones needed for both proper ecdysis and dauer formation [49, 50], mutations in cholesterol metabolism and steroid signaling would be expected among yet-undescribed Daf-c mutants.

To identify the key molecular factors controlling dauer-specific phenotypes, we examined the development and behavior of *P. pacificus* mutants that form DL constitutively. In a previous study, a forward genetic screen in *P. pacificus* yielded four dauer constitutive Daf-c alleles, *dfc* (<u>d</u>auer-formation <u>c</u>onstitutive), which can be rescued by exogenous Δ7-dafachronic acid [8, 51]. One of the strongest alleles, *Ppa-dfc-1(tu391),* has been positionally mapped, but the genetic lesion remains unknown. We therefore isolated a new Daf-c allele, *csu60*, and following positional mapping and whole genome sequencing, found that *csu60* is a null allele of the sole hydroxysteroid dehydrogenase in *P. pacificus*, an enzyme responsible for the production of dafachronic acids. For both Daf-c alleles, the DL and adults have olfactory behavior resembling wild-type starvation-induced DL, suggesting that the genes which control dauer entry also normally repress host-seeking olfactory behavior in adults.

## Results

### *P. pacificus* Daf-c mutants have an exsheathment phenotype

*P. pacificus* dauer larvae (DL) can be induced by starvation, crowding, or high temperature (≥25°C). We characterized two incompletely penetrant, recessive, non-allelic dauer formation-constitutive (Daf-c) mutants at 20°C, *Ppa-dfc-1(tu391)* and *csu60*. Like several species of parasitic nematodes, most wild-type *P. pacificus* DL are ensheathed (63%), where the sheath is the apolysed but unecdysed cuticle of the J2 larvae [52]. Strikingly, while the ensheathed wild-type DL are motile, both Daf-c alleles exhibited an unusual ensheathment-related defect in which live worms are trapped and immobilized inside an old cuticle (Fig. 1A-C) [53]. Aside from the loosened cuticle, most “incarcerated” larvae in *tu391* and *csu60* are indistinguishable from DL produced by the wild-type reference strain PS312 under starvation condition: they are characterized by radial shrinkage of the body, a darkly hued gut, and a buccal cap covering the mouth. In entomopathogenic nematodes from the genera *Steinernema* and *Heterorhabditis*, ensheathment appears to confer increased ability to resist desiccation, while exsheathment leads to increased motility and is the first step of the host infection process [53, 54]. Our results reveal that in *P. pacificus*, exsheathment is genetically coupled with dauer entry.

**Figure 1.**
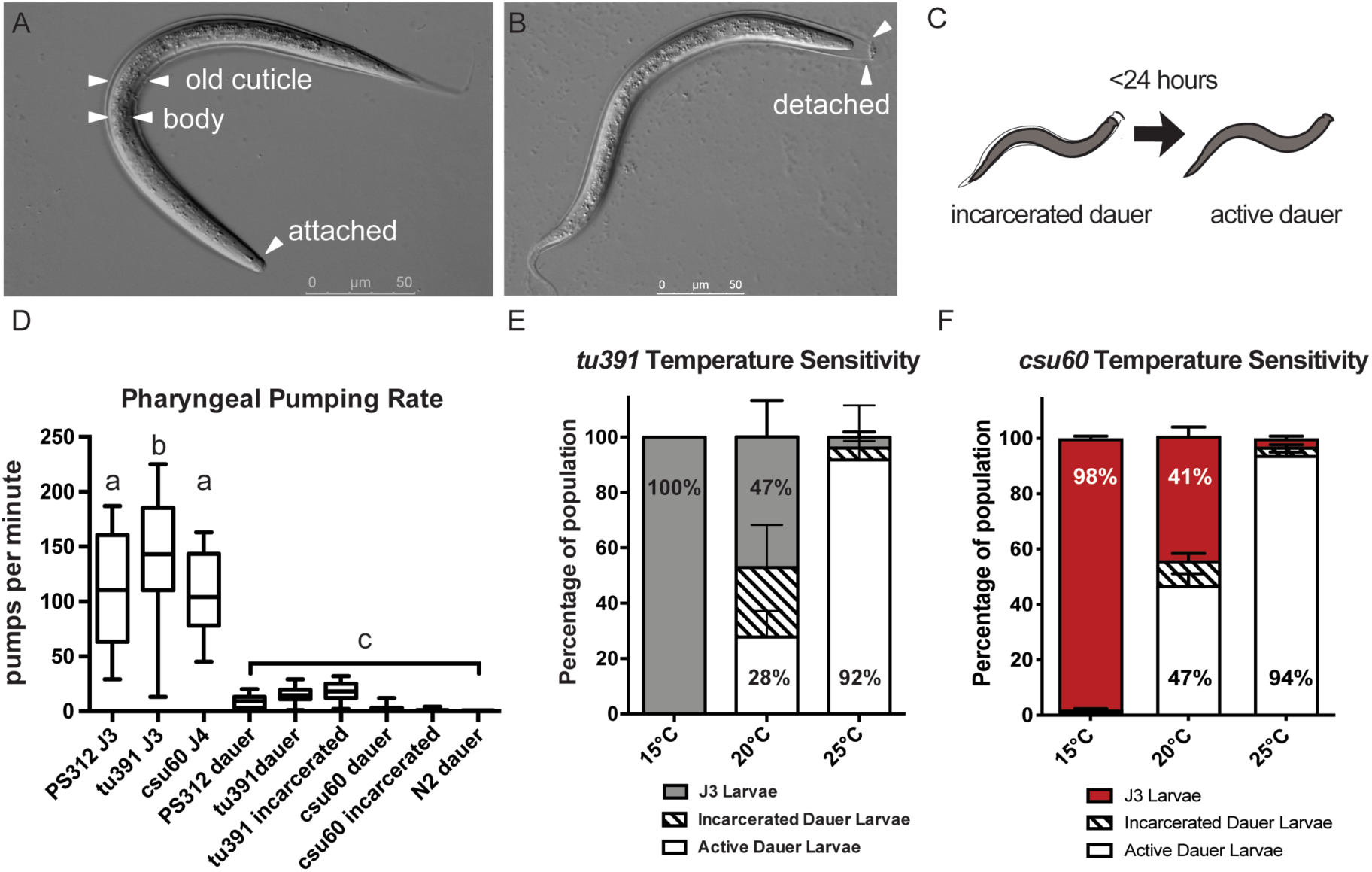
Incarcerated larvae are temperature-dependent constitutive dauer larvae. Ten young adult hermaphrodites on food were transferred to different temperatures for 5 days, and their progeny were then scored for dauer-equivalent J3 stages. **(A)** Ensheathed wild-type PS312 dauer larvae. The effete J2 cuticle surrounding the body is visible. **(B)** In an incarcerated *tu391* dauer larva, the J2 cuticle is not attached at the head in nearly half of *tu391* constitutive dauer larvae. **(C)** The illustration depicts that the incarceration is transient. **(D)** Incarcerated dauer larvae (n_tu391_=22; n_csu60_=30) exhibit pharyngeal pumping rates similar to active mutant dauer (n_tu391_=22; n_csu60_=13) and wild-type *P. pacificus* and *C. elegans* dauer (n_Ppa_=24; n_Cel_ =18), which are significantly different from that of the J3 and J4 larval stages (PS312 J3 n=32; *tu391* J3 n=40; *csu60* J4 n=13) (letters denote significant differences at *P*<0.001, One-way ANOVA, Tukey’s Multiple Comparisons test). **(E)** The proportions of *tu391* J3 larvae, active dauer larvae, and dauer larvae differ at 15°C (n=577), 20°C (n=2091), and 25°C (n=1622). Incarcerated dauer larvae are 25% of all third-stage larvae larval stage animals at 20°C. Wild-type PS312 are 100% J3 at all temperatures. **(F)** The proportions of *csu60* J3 larvae, active dauer larvae, and incarcerated dauer larvae differ at 15°C (n=1250), 20°C (n=1891), and 25°C (n=1466). Incarcerated dauer larvae are 9% of all third-stage larvae larval stage animals at 20°C. Additional details are found <u>in</u> **Table 1**.

**Table 1.** Phenotype frequency in Daf-c mutants

| Strain | Daf-c DL<sup>a</sup><br>±SEM (%)<br>@25°C | N | Active DL<br>±SEM (%)<br>@20°C | Mlt defect<br>±SEM (%)<br>@20°C | N | Daf-c DL<sup>a</sup><br>±SEM (%)@15°C | N |
| --- | --- | --- | --- | --- | --- | --- | --- |
| **Wild-type PS312** | 0 | 512(3) | 0 | 0 | 482(3) | ND |  |
| *hsd-2(csu60)* | 99 ±0.4 | 1577(5) | 47 ±2 | 9 ±1.2<sup>b</sup> | 1891(5) | 2 ±0.4 | 1250(5) |
| *hsd-2(csu60); daf-12(tu389)* | 0 | 1172 (6) | 0 | 3.7±3.4<sup>c</sup> | 808(6) | ND |  |
| *daf-12(tu389)* | 0 | 1250(7) | 0 | 0 | 950(7) | ND |  |
| *hsd-2(csu60); csuEx54[Ppa-*<br>*hsd-2p::hsd-2 cDNA]* | ND |  | 11 ±2.5 | 0 | 2388(6) | ND |  |
| *hsd-2(csu60) + Δ7-DA* | ND |  | 0 | 0 | 284(3) | ND |  |
| *dfc-1(tu391)* | 96 ±0.9 | 1520(7) | 28 ±3.5 | 25 ±2.6<sup>b</sup> | 1742(11) | 0 | 577(7) |
| *dfc-1(tu391); daf-12(tu389)* | 0 | 1005(7) | 0 | 0 | 1003(7) | ND |  |
| *dfc-1(tu391) + Δ7-DA* | 0 | 426(4) | ND | ND |  | ND |  |

Values based on the progeny of 10 gravid hermaphrodites scored 5 or 6 days later.
<sup>a</sup>Includes both active and exsheathment defective incarcerated dauer larvae formed under reproductive conditions among all J3-equivalent larvae.
<sup>b</sup>Transient exsheathment defective Daf-c dauers; no molting defect was found in non-dauers.
<sup>c</sup>Transient molting defect in J3-J4 or J4-Adult molts.
N: sample size with the number of experiments is given in parentheses.
ND: not determined
SEM: standard error of the mean

While most of the *csu60* incarcerated larvae resemble DL, some animals show less radial constriction than wild-type dauers. To confirm that all incarcerated worms are exclusively in the dauer stage, we measured their pharyngeal pumping frequency. We found that the incarcerated *tu391* and *csu60* larvae display pumping rates significantly lower than the J3 and J4 stages of wild-type and mutant animals, but indistinguishable from their respective non-incarcerated (active) DL as well as wild-type *P. pacificus* and *C. elegans* DL (Fig. 1D). We isolated *tu391* and *csu60* J3 and J4 larvae onto fresh OP50 to determine if they develop into incarcerated larvae but did not observe incarcerated worms. Furthermore, we transferred mutant *tu391* J2 larvae onto fresh *E. coli* OP50 and found that J2 larvae formed J3 larvae, active DL, or incarcerated DL, suggesting that the molting defect in *tu391* and *csu60* is exclusive to the J2-DL transition.

To determine if the incarcerated DL is a permanent or transient phenotype, we picked incarcerated *tu391* and *csu60* DL onto fresh OP50 plates. Because transferring the incarcerated DL may trigger the release the trapped DL, we also isolated the incarcerated *tu391* DL by removing all other worms from the plate rather than picking the incarcerated *tu391* DL. Surprisingly, all ‘disturbed’ and ‘undisturbed’ incarcerated *tu391* and *csu60* DL became free and active within 24 hours, regardless of whether they were maintained at 20°C or 25°C. Thus, the incarcerated DL is a transient dauer-specific phenotype that effectively results in exsheathed *tu391* and *csu60* DL, which highlights the presence of a genetic regulator in *P. pacificus* that coordinates DL formation with proper cuticle ensheathment. Despite superficial resemblance between ensheathed wild-type DL and Daf-c DL with the exsheathment defect, only the Daf-c mutations result in immobilized DL, suggesting that the same genes regulate dauer entry and J2-DL cuticle ensheathment in *P. pacificus*.

Because temperature modulates the dauer decision [55, 56], we wondered if the constitutive dauer formation phenotypes and the dauer-specific molting defects are temperature-dependent. We placed ten young adult hermaphrodites on OP50 at different temperatures for 5 days and scored for dauer-equivalent stages. We found that for *tu391,* lower temperature (15°C) completely suppresses the Daf-c phenotype, while higher temperature (25°C) significantly enhances the Daf-c phenotype but dramatically suppresses the dauer incarceration defect (Fig. 1E). 25°C promotes the highest proportion of free DL (92% of third-stage larvae). For *csu60,* lower temperature also suppresses the Daf-c phenotype, while 20°C promotes the highest proportion of incarcerated DL (9%) (Fig. 1F). Similar to *tu391* and other Daf-c alleles in *C. elegans*, free DL formation in *csu60* is highest at 25°C (94%) compared to 20°C (47% of third-stage larvae) [55, 57]. In contrast, wild-type cultures had 100% J3 larvae at 20°C and 25°C (Table 1). Thus, increasing the temperature resulted in higher DL formation in both alleles, while exsheathment defects are most prevalent at 20°C. Higher temperature may suppress the initiation of the exsheathment defect, or it may shorten the duration of the incarceration. Together, these findings indicate that the Daf-c mutants in *C. elegans* and *P. pacificus* both promote dauer formation at higher temperature, but the Daf-c alleles in *P. pacificus* show an additional cuticle exsheathment phenotype at 20°C.

### Chemosensory sensilla of Daf-c mutants

In *C. elegans*, a bilaterally symmetrical pair of anterior sensory organs known as the amphid sensilla mediates chemosensory responses to water-soluble compounds and volatile odors. To determine if the Daf-c mutations affect the formation or function of the amphid neurons, we used the lipophilic live dye DiI to stain a subset of the amphid neurons directly exposed to the external environment. DiI stains a stereotypic set of neurons by retrograde transport that appears to have a modest degree of conservation across various nematode species, including *P. pacificus* and *C. elegans* [58–60]. Moreover, several abnormal dye-filling-defective mutants in *C. elegans,* known as the Dyf mutants, also have dauer formation phenotypes; most are dauer-formation defective (Daf-d), while a few are Daf-c [61–64].

We found that three of the twelve pairs of amphid neurons consistently stain with DiI in wild-type *P. pacificus* J2, dauer, and J4 larvae, as well as young adults. Based on the recent amphid neuron homology assignments between *P. pacificus* and *C. elegans* [60], these amphid neurons correspond to the ASK, ADL, and ASH neuronal homologs. Interestingly, the posterior phasmid neurons that dye-fill in all developmental stages in *C. elegans* only show DiI fluorescence in the *P. pacificus* DL. We found that the majority of *tu391* J2 larvae (65%) and J4/adults (97%) stain very weakly in these three pairs of amphid neurons, suggesting possible defects in the differentiation or function of either the amphid neurons or the amphid sheath glial cells that support them (Fig. 2A-D). In addition, 22% of *tu391* DL showed a noticeable decrease in DiI uptake compared to 12% of wild-type DL (Fig. 2E-F). Using the amphid sheath cell marker *Ppa-daf-6p::rfp* [60], we found superficially wild-type expression in *tu391* adults; thus, the defect in dye update is likely due to amphid neurons rather than glia (SI Fig. 1). In contrast to *tu391*, most *csu60* adults (88%) take up DiI close to the rate of wildtype (97%)(SI Fig. 2). The reduction of DiI uptake was the most severe in the *tu391* J2 larvae and adults (weak staining in 97% *tu391* vs. 3% wild-type adults), but dramatically less severe in *tu391* DL (weak staining in only 22% *tu391* vs. 12% wild-type DL). These results suggest that: (1) the amphid sensilla are remodeled during dauer development, such that *tu391* DL are not as dye-filling defective as *tu391* J2 and adults are; (2) the two Daf-c alleles have pleiotropic phenotypes related to distinct aspects of dauer entry in *P. pacificus*.

**Figure 2.**
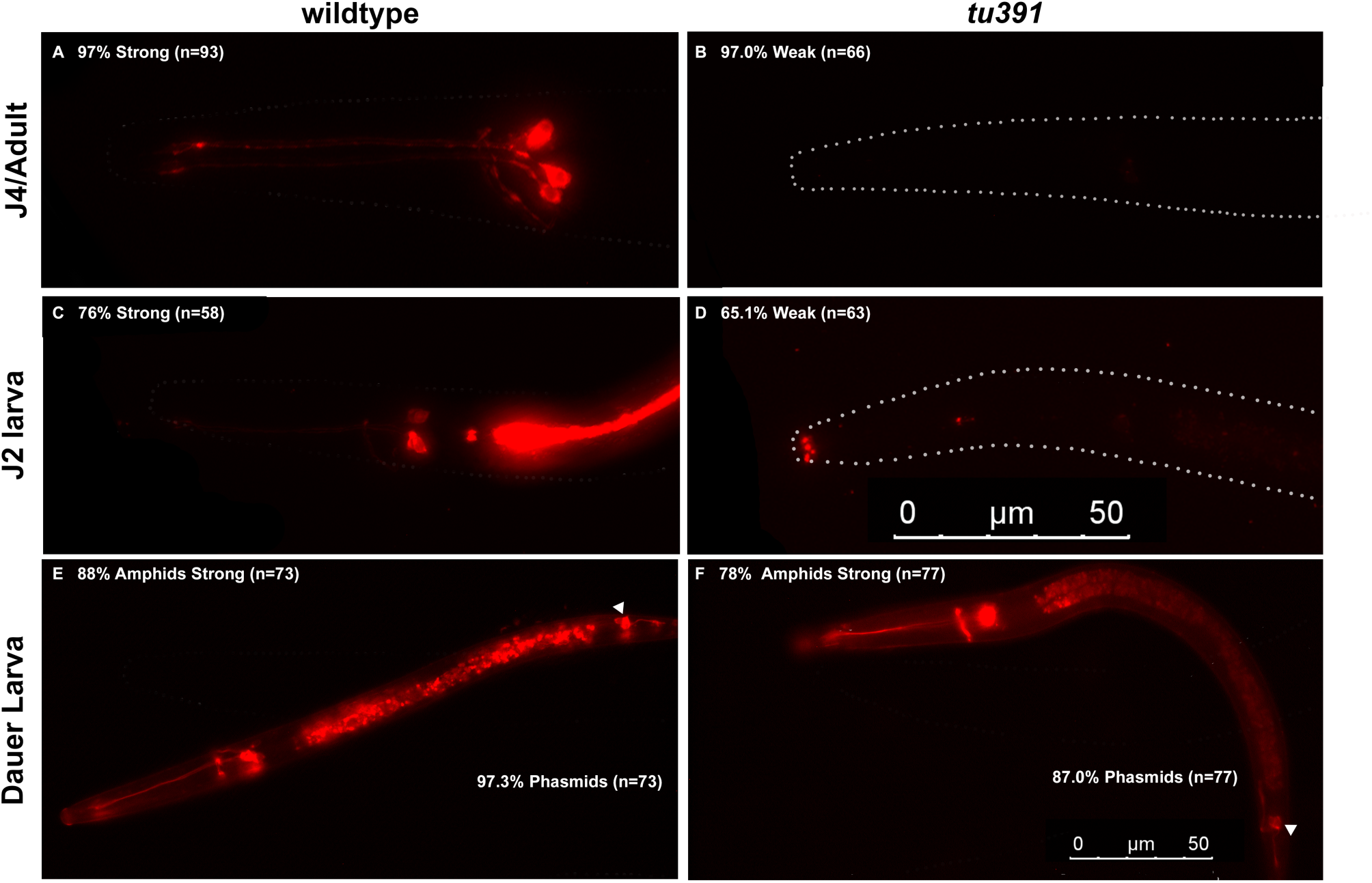
*tu391* is defective in DiI uptake. (A-D) Adult, J4, and J2 *tu391* larvae show significantly weaker staining in the amphid neurons when compared to wild-type PS312 animals. **(E-F)** However, the *tu391* dauer larvae take up dye at a level comparable to wild type, including in the posterior phasmid neurons that take up dye only in the DL stage (triangle). Worms were exposed to 6.7 nM DiI in M9 for 3 hours and scored categorically based on dye intensity and the number of visible neurons. 0.1% SDS with M9 was used for staining dauer larvae. Max projections of stacked images, exposure 200 ms (A, B) or 524 ms (C-F). Anterior is left and dorsal is up. The 50 µm scale bar in (D) represents (A-D), and the scale bar in (F) represents (E-F).

### Daf-c mutations alter adult chemotaxis response

Since most chemotaxis responses in non-parasitic nematodes have been based on assays conducted with young adults [23-25,39], we sought to use the high proportion of active Daf-c DL to examine potential behavioral changes between young adults and DL. Performing chemotaxis assays on wild-type *P. pacificus* DL is difficult, not only due to the paucity of dauers induced by starvation but also because the cuticle surface of *P. pacificus* DL is covered by the long-chain polyunsaturated wax ester Nematoil [35], which retards movement on the standard chemotaxis medium [24, 39]. Therefore, we adopted a modified agar medium containing the detergent Tween, which promotes dispersal [65]. To determine if Daf-c DL behave similar to wild-type DL, we tested DL of wild type, *tu391,* and *csu60* with a panel of odorants that are known attractants for *P. pacificus* adults. We found that the DL of wild type and *csu60*, and to a lesser extent *tu391*, exhibited strong attraction to the Oriental beetle pheromone ZTDO and the fly aggregation pheromone methyl myristate (Fig. 3A). Given this wild-type chemosensory response by the Daf-c DL, we expanded the survey of DL chemosensation to the other insect pheromones E-TDA, hexadecanal, and ß-caryophellene but found that the Daf-c DL show no response to these odorants (SI Fig. 3) [39,66,67]. We also examined the responses of Daf-c DL to a panel of *C. elegans* attractants: 2,3-butanedione (also called diacetyl), 2,3-pentanedione, and isoamyl alcohol (IAA) [24, 39]. We found that the DL do not respond to these chemicals; *P. pacificus* adults also do not respond (SI Fig. 3). Thus, the Daf-c DL exhibited near wild-type responses to two known host-associated odors but have a narrower overall odor response profile compared to wild-type adults. It remains to be determined the degree of concordance between wild-type and Daf-c DL chemosensory profiles.

**Figure 3.**
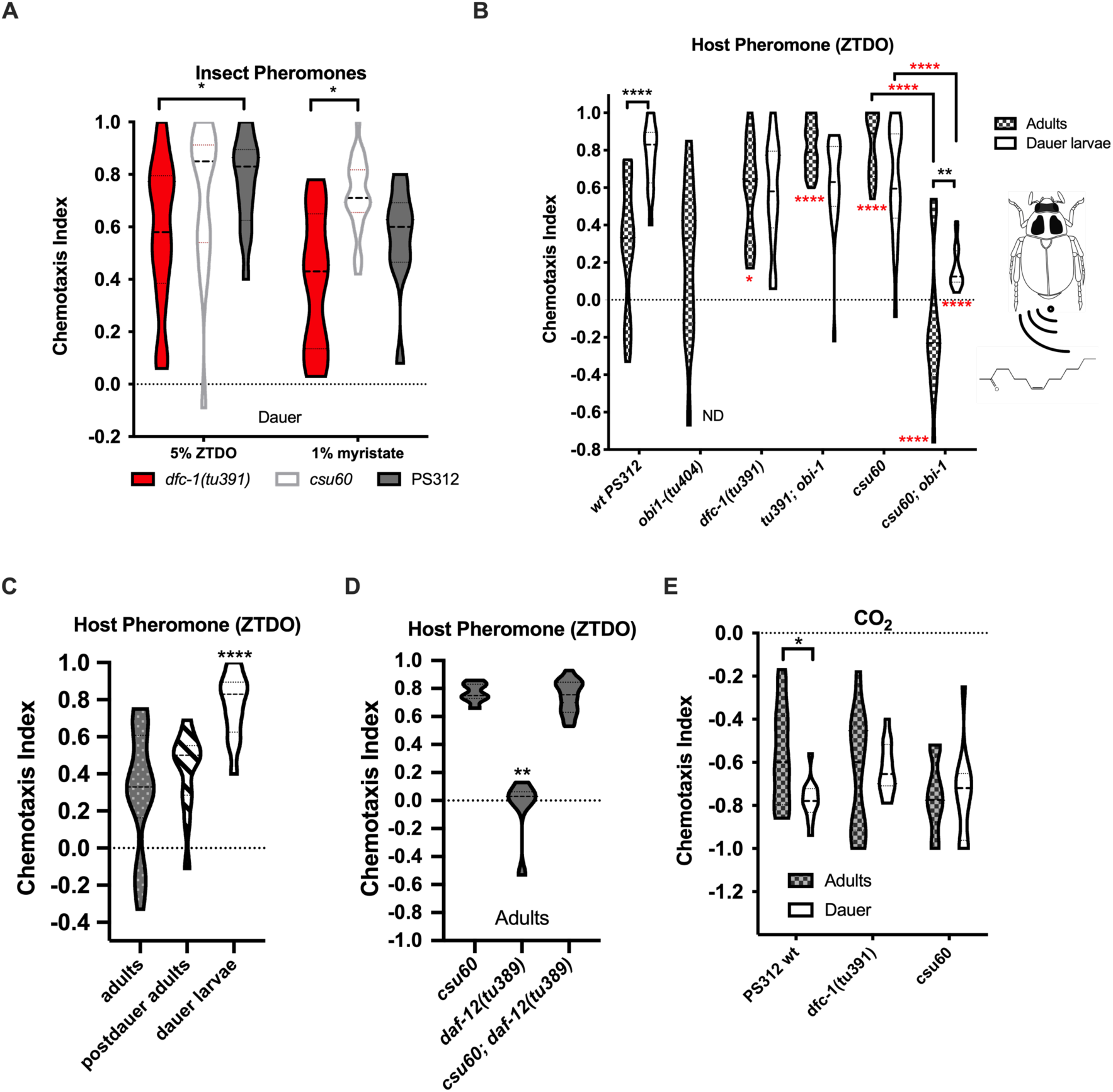
Olfactory responses of *P. pacificus* dauer larvae. (A) Dauer larvae of wild-type and Daf-c mutants show robust attraction to 5% Oriental beetle pheromone z-(7)-tetradecen-2-one (ZTDO) and 1% house fly aggregation pheromone methyl myristate. Two-way ANOVA with Dunnett’s multiple comparisons test to wildtype. **(B)** Daf-c mutations enhance adult attraction to the host beetle pheromone ZTDO in *tu391* and *csu60*, while the loss of *obi-1* significantly reduces attraction *csu60* attraction to ZTDO. Two-way ANOVA for differences by genotype with Tukey’s multiple comparisons test in red asterisks; asterisks below show differences to PS312. Black asterisks denote differences between adults and dauer by Sidak’s test. **(C)** Post-dauer adults show a similar response to ZTDO as the never-dauer adults. One-way ANOVA against adults with Dunnett’s post-hoc test. **(D)** The steroid hormone receptor encoded by *Ppa-daf-12* is not required for the enhanced attraction to ZTDO in adult *csu60*. One-way ANOVA against *csu60* with Kruskal-Wallis test. **(E)** The response to carbon dioxide differs slightly between wild-type adults and dauers, but does not differ between adults and dauer larvae in the Daf-c mutants. Two-way ANOVA between adults and dauer with Sidak’s test. For each category, 12-15 assays were performed over three trials. Violin plots show median with quartiles. \**P*<0.05; \*\**P*<0.01; \*\*\*\**P*<0.0001.

To better define the genetic pathways that are responsible for the chemosensory differences between DL and adults, we focused on the responses of the Daf-c mutants to the Oriental beetle pheromone ZTDO [68]. We observed that wild-type PS312 showed the strongest difference between the DL and adults, with an almost threefold stronger attraction in DL than in adults (Fig. 3B). We also observed significantly stronger attraction by the adults of both Daf-c alleles, such that there were no significant differences between the responses of DL and adult stages. The *csu60* adults were significantly more attracted to ZTDO than both wild-type and *tu391* adults, and despite the dye-filling defect in *tu391* being more pronounced in adults compared to DL, the attraction to ZTDO is stronger in *tu391* adults than in wild-type adults. Because the passage through the dauer stage may introduce lasting post-dauer changes that alter odorant receptor expression and hence modulate behavior [7,69,70], we also examined the ZTDO response of wild-type post-dauer adults. We found that dauer passage did not result in enhanced ZTDO response in adults (Fig. 3C). Thus, the enhanced ZTDO attraction in Daf-c adults was not due to developmental experience, but rather the heterochronic activation of a dauer-specific neuronal development.

To determine how the Daf-c mutants interact with chemosensory mutants mediating the ZTDO response, we examined the chemotaxis behavior of the Daf-c mutants in the loss-of-function *obi-1(tu404)* background that is defective for its cGMP-dependent responses to ZTDO [68]. We found that the *obi-1* background did not affect ZTDO attraction in both *tu391* adults and DL (Fig. 3B). In contrast, the loss of *obi-1* significantly reduced the ZTDO response in *csu60* adults as well as DL, from strong enhanced attraction to weak repulsion and neutrality, respectively (Fig. 3B). We were unable to perform chemotaxis assays on the DL of *obi-1(tu404)* mutants due to its high DL mortality rate, especially in the presence of ZTDO [68]. Interestingly, while *Ppa-daf-12* is epistatic to *csu60* for dauer formation, *csu60*; *daf-12(tu389)* double mutant adults still exhibited the *csu60* enhanced attraction to ZTDO (Fig. 3D). Thus, the regulation of chemotaxis responses can be uncoupled from dauer formation and requires a steroid hormone that does not act through a DAF-12 homolog. Taken together, our results suggest that *tu391* acts downstream of or in parallel with *obi-1*, while *csu60* likely acts upstream of *obi-1* and independent of *daf-12* to mediate the enhanced pheromone attraction of young adults.

Since host odor attraction in IJs of entomopathogenic nematodes is enhanced by carbon dioxide (CO_2_) [21], we next tested the chemotaxis response of *P. pacificus* DL to CO_2_. In *C. elegans*, well-fed adults are repelled by CO_2_ while DL are attracted to the same concentration [20, 21]. In *P. pacificus* well-fed adults, CO_2_ also elicits a strong avoidance response which, as in *C. elegans*, is mediated by a pair of internal gas sensing neurons known as the BAG neurons [20]. To determine if CO_2_ response valence also differs between *P. pacificus* DL and adults, we examined the response to CO_2_ in Daf-c DL and adults using the modified chemotaxis assay. We found that wild-type *P. pacificus* DL exhibited a slightly stronger avoidance response than adults, while both Daf-c DL and adults showed equally robust repulsion by CO_2_ (Fig. 3E). Thus, similar to the enhanced response to the beetle host pheromone, wild-type *P. pacificus* DL displayed a slightly enhanced negative response to CO_2_, not a valence change. Interestingly, iL3s of the mammalian parasites *S. stercoralis*, *S. ratti, and Nippostrongylus brasiliensis* are also repelled by CO_2_ [22]. It is currently unclear how the CO_2_ avoidance response of *P. pacificus* DL, but not *C. elegans* DL, factors into the different life strategies of these two species in the wild.

Finally, to determine if genes that affect dauer development also modulate olfactory behavior in other nematodes species, we also compared the odor response profiles of young adults vs. DL in *C. elegans* [71], including the odor response profiles of two well-studied Daf-c mutants, *daf-2(e1370)* and *daf-7(e1372, ok3125)* [56, 72]. We found that DL of wild-type N2 and both the Daf-c mutants showed weaker odorant responses than adults. Specifically, wild-type DL showed lower attraction than adults to 2,3-butanedione, 2,3-pentanedione, and IAA, consistent with the results of a previous study that included 2,3-butanedione [27](Fig. 4A-C). Similar to the finding in *P. pacificus*, we also found dauer passage did not result in enhanced IAA or 2,3-pentanedione attraction in post-dauer adults (SI Fig. 4). Most unexpectedly, the response to IAA exhibited valence changes between adults and DL: it changed from attractive to neutral in wild-type animals, and from attractive to repulsive in both *daf-7* mutants. To evaluate if the avoidance response to IAA is due to hypersensitivity in the *daf-7* DL, we also tested a 10-fold lower IAA concentration (0.1%) and found that IAA attraction was still significantly reduced in both *daf-7* alleles compared to wild type (Fig. 4D). Thus, *C. elegans* wild-type and *daf-2* DL showed decreased attraction compared to adults to all three odorants tested, but more dramatically, *daf-7* alleles showed a valence change by accentuating the avoidance response of DL to IAA, suggesting that the TGF-ß pathway is involved in both dauer regulation and IAA sensing.

**Figure 4.**
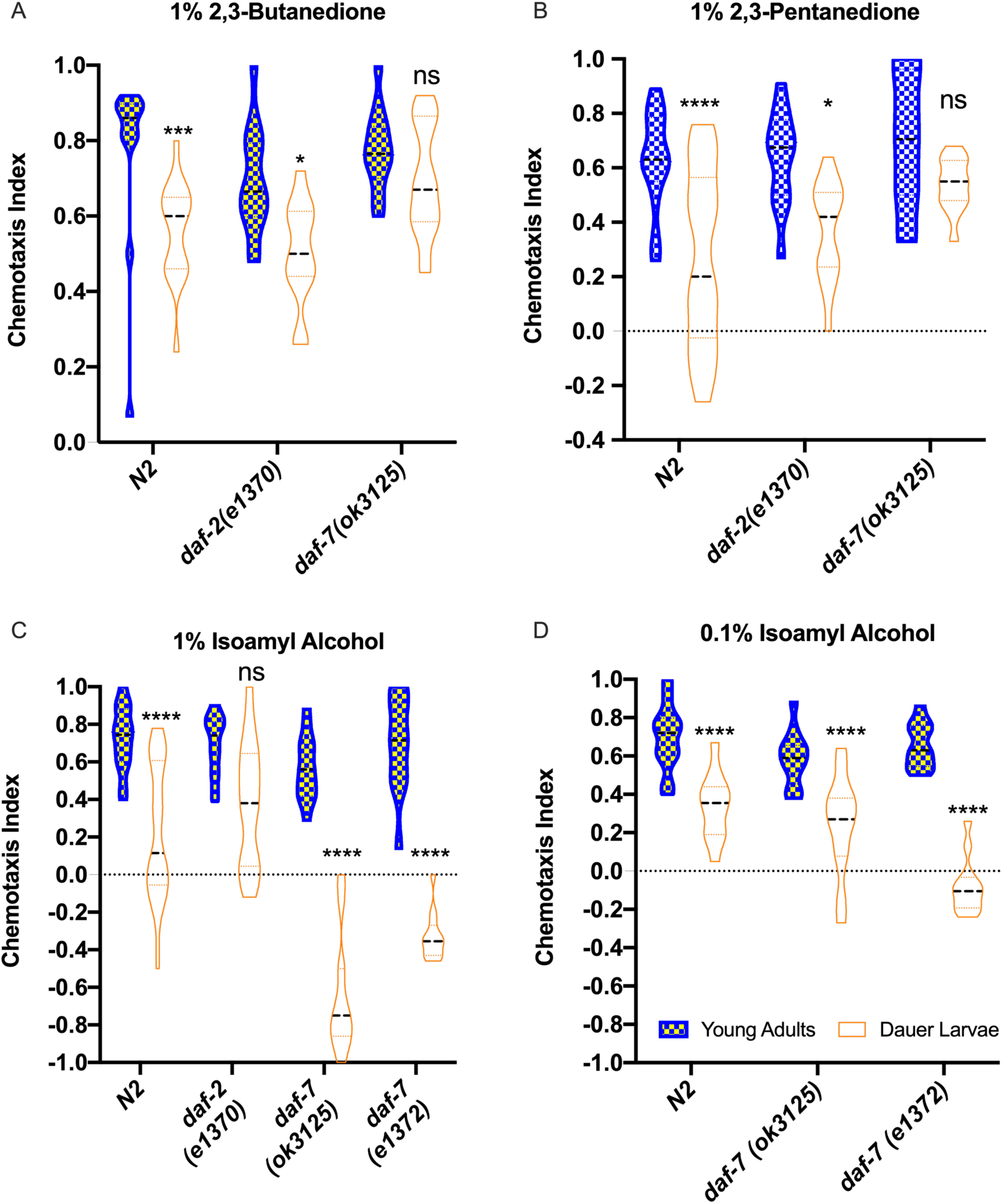
Olfactory responses of *C. elegans* dauer larvae. Chemotaxis between adults and dauer larvae differ significantly in response to **(A)** 1% 2,3-butanedione, **(B)** 1% 2,3-pentanedione, and **(C-D)** 1% and 0.1% isoamyl alcohol in wild-type N2 and the Daf-c mutants *daf-2* and *daf-7.* For A and C: One-way ANOVA between adults and dauer with Kruskal-Wallis test. For B and D: Two-way ANOVA between adults and dauer with Sidak’s multiple comparisons test. \**P*<0.05, \*\**P*<0.01, \*\*\*\**P*<0.0001, ns: not significant. For each category, 12 assays were performed over >3 trials. Violin plots show medians with quartiles.

### Molecular cloning of a Daf-c mutant

In *C. elegans*, the binding of the nuclear hormone receptor DAF-12 to the steroid hormones Δ4-dafachronic acids and Δ7-dafachronic acids (DAs) restrains L2 larvae from undergoing the dauer fate [51]. To determine if *csu60* or *tu391* act upstream of *Ppa-daf-12*, we tested a synthetic DA for its ability to ameliorate the Daf-c phenotype. Previous work has shown that the constitutive dauer formation phenotype of *dfc-1(tu391)* is rescued by feeding the worms *E. coli* OP50 containing 7.5 µM Δ7-dafachronic acid (Δ7-DA) [8]. Δ7-DA has been shown to prevent dauer formation in both *P. pacificus* and the ruminant parasite *Strongyloides papillosus* [8]. We found that both the dauer-specific exsheathment defect as well as the constitutive DL phenotype of *dfc-1(tu391)* and *csu60* were rescued by the availability of Δ7-DA in the food *ad libitum* (Fig. 5A). These results suggest that neither mutant produces the *P. pacificus* dafachronic acid that is functionally equivalent to the dauer-suppressing Δ7-DA.

**Figure 5.**
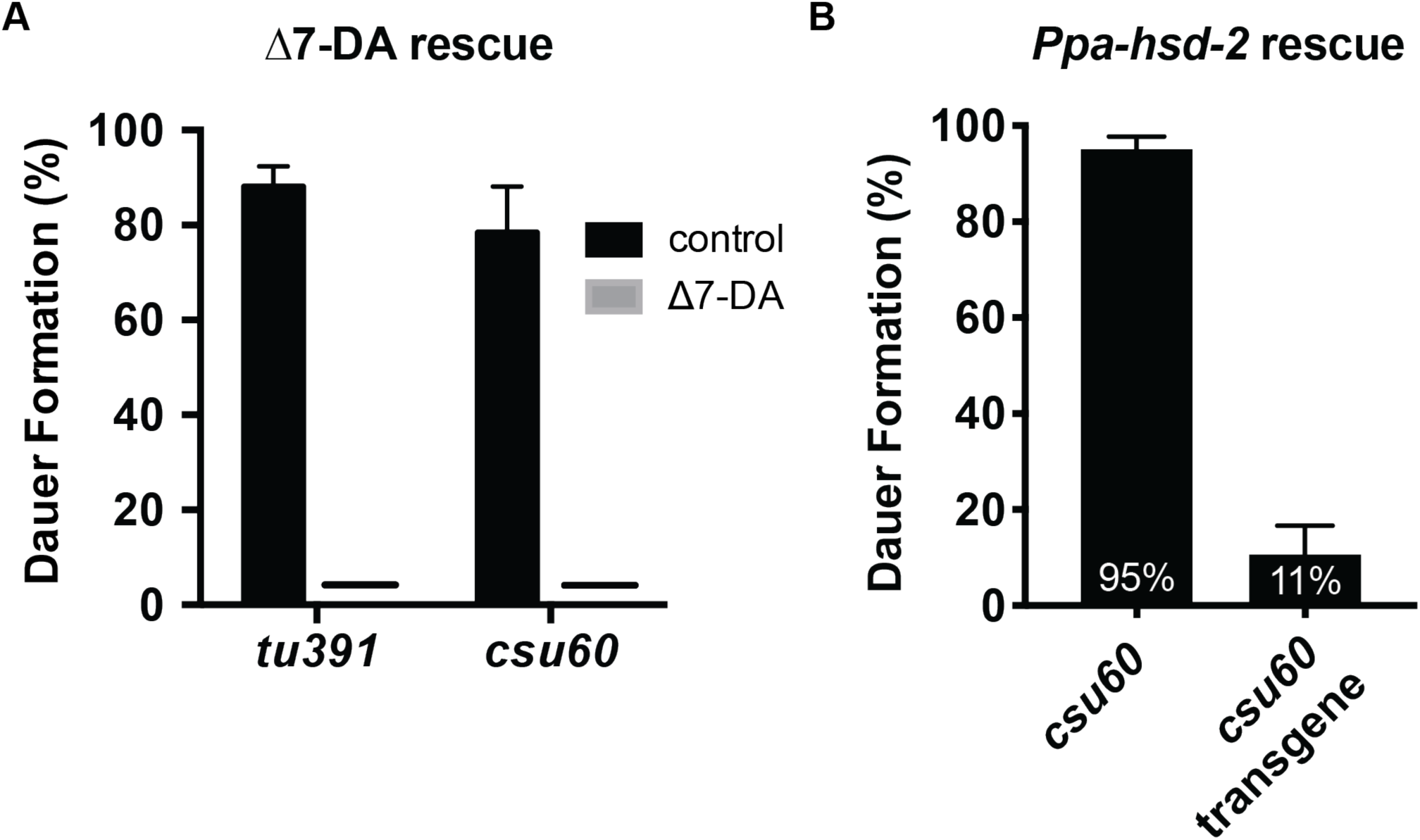
Hormonal and transgenic rescue of Daf-c mutants. (A) The constitutive dauer phenotype in both Daf-c mutant alleles is completely rescued by exogenous Δ7-dafachronic acid but not by the ethanol control. (T-test between mock and DA treated. *P*<0.0001. Sample size: *tu391* control = 435; *tu391* treated = 426; *csu60* control = 210; *csu60* treated = 284). **(B)** A transgene carrying the *Ppa-hsd-2* promoter driving its own cDNA strongly rescues the constitutive dauer defect in *csu60* (T-test between transgenic and non-transgenic populations. *P*<0.0001; Sample size: *csu60* = 2246; *csu60* segregating for the rescue transgene = 2388). Dauer formation is the percentage of DL in all third-stage larvae or older.

To further confirm this genetic pathway for dauer regulation by epistasis analysis, we constructed *csu60; daf-12(tu389)* double mutants and found that they resembled the dauer formation defective phenotype of *daf-12(tu389)* [8] (Table 1), consistent with the expected position of *csu60* acting upstream of the nuclear hormone receptor (see Methods). Similarly, the *dfc-1(tu391); daf-12(tu389)* double mutants completely masked the Daf-c phenotypes of *tu391*, placing *tu391* upstream of *daf-12* for the regulation of dauer formation. Hence, *daf-12* is epistatic to both *csu60* as well as *tu391*. A small percentage (3.7%) of the *csu60; daf-12(tu389)* double mutants exhibited an ecdysis defect in the molts between late larval stages (J3-J4; J4-adult), which is a synergistic phenotype not found in either single mutant (Table 1). These results suggest that both mutations disrupt the steroid biosynthetic and signaling pathways critical for dauer formation and possibly molting.

Using the latest genome assembly of *P. pacificus* El Paco, we performed whole genome sequencing of *csu60* and identified a ∼95 kb deletion on Chromosome II that contains 11 predicted protein-coding genes. Among these candidates, the predicted protein sequence of PPA10139 is orthologous to the 3β-hydroxysteroid dehydrogenase/Δ5-Δ4 isomerases (3β-HSDs) that are key steroidogenic enzymes in vertebrates (GO:0003854). PPA10139 shares 89% similarity with *C. elegans* HSD-2, which encodes one of three hydroxysteroid dehydrogenases in *C. elegans* involved in the biosynthesis of dafachronic acids [73]. Consistent with its potential role in repressing dauer formation, PPA10139 transcripts are significantly lower in DL than in the J2, J3, J4, and adult stages (gene prediction Contig6-snapTAU.237) [74]. To confirm that the deletion of *Ppa-hsd* is solely responsible for the Daf-c phenotype of *csu60*, we made a construct containing a ∼2 kb promoter of PPA10139 driving its full-length cDNA to rescue the *csu60* deletion by transgene complementation. Compared to the *csu60* mutant, the *csu60* extrachromosomal transgenic rescue line showed dramatically lower constitutive dauer formation (Fig. 5B). Thus, both the constitutive dauer and exsheathment defects are caused by the genomic deletion containing the only predicted *P. pacificus hsd* homolog.

When we investigated the possible evolutionary history of *hsd* genes in nematodes, we found only one putative ortholog of HSD-2 in each of the completed genomes of the human parasites *Brugia malayi* and *Onchocerca volvulus*, indicating that a single HSD is likely the ancestral state, whereas the *C. elegans* genome contains three copies (Fig. 6A). Each *hsd* gene in *C. elegans* differs by the number of transmembrane domains: *hsd-1* encodes for two predicted domains, *hsd-2* encodes for one domain, while *hsd-3* does not encode for any predicted transmembrane domains (as predicted by SMART, which utilizes TMHMM to predict transmembrane domains). Although the overall protein sequence of PPA10139 shares the highest similarity to *C. elegans* HSD-2, the two transmembrane domains predicted for *Ppa-*HSD-2 argues for more functional similarity to *C. elegans* HSD-1 (Fig. 6B).

**Figure 6.**
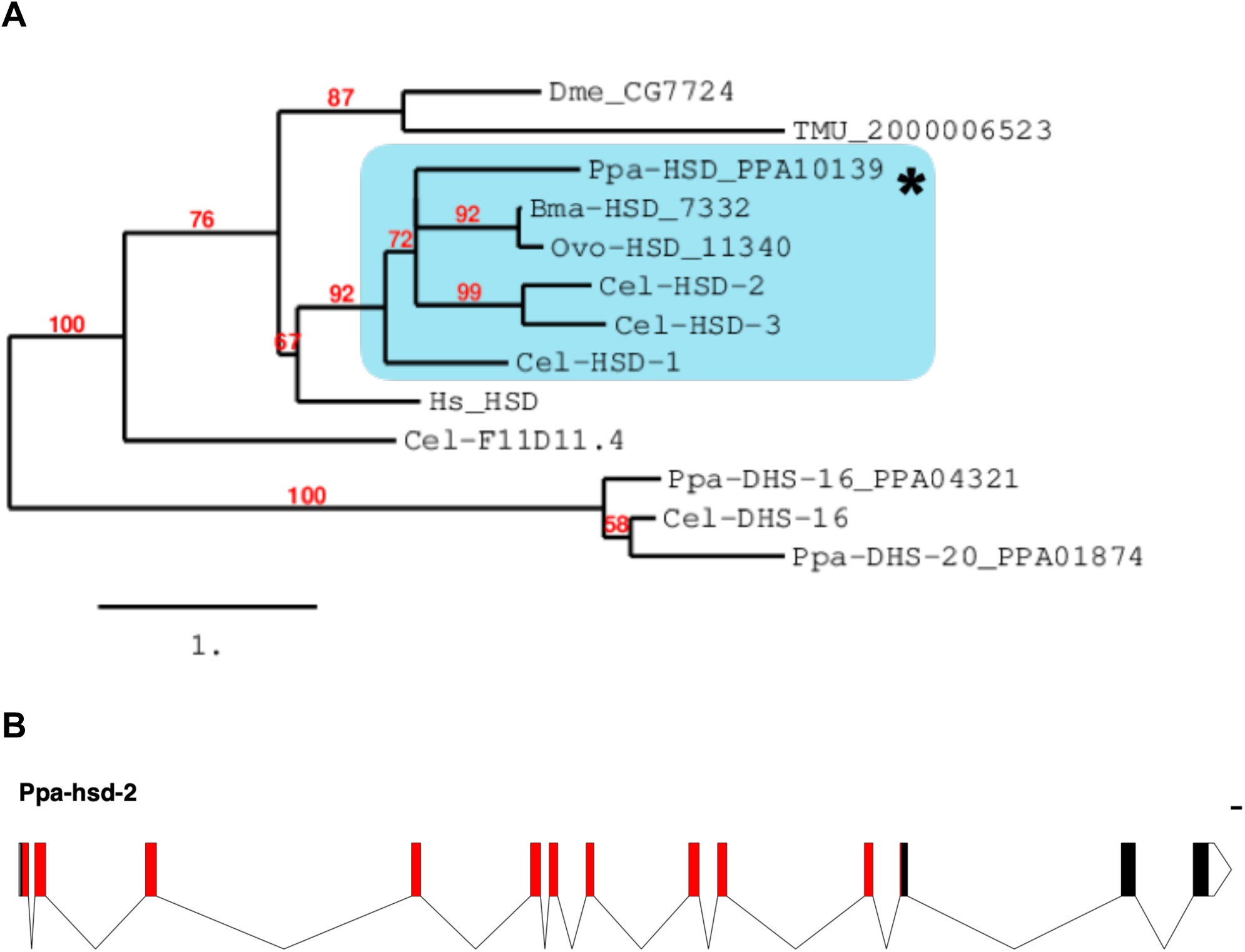
*Ppa-hsd-2* phylogeny and gene annotation. (A) Orthology assignment for *Ppa-*HSD-2 (PPA10139). Maximum Likelihood phylogeny trees were generated by PhyML (phylogeny.fr). Midpoint rooting was used and only branch support ≥50% is shown. Protein sequences for *P. pacificus* are “Ppa-” (orthology same as Wormbase.org) and for *C. elegans* “Cel-“. Sequences from additional species are *Drosophila melanogaster* (Dme), *Onchocerca volvulus* (Ovo), *Brugia malayi* (Bma), *Trichuris muris* (Tme), and *Homo sapiens* (Hs). Dehydrogenases DHS-16 from *C. elegans* along with DHS-16 and DHS-20 from *P. pacificus* are closest to the HSDs and are used as an outgroup. **(B)** *Ppa-hsd-2* gene annotation (PPA10139). Red denotes the predicted 3 beta-hydroxysteroid dehydrogenase protein family spanning amino acid residues 6-274 (exons 1-11). Gene annotation was created using the Exon-Intron Graphic Maker on wormweb.org, and the SMART webtool was used to predict protein domains (smart.embl-heidelberg.de Letunic *et al*., 2014 and 2017). Unfilled blocks are 5’ and 3’ UTRs. Scale bar: 100 bp.

We hypothesized that if *Ppa-*HSD-2 is the only hydroxysteroid dehydrogenase in *P. pacificus* involved in dafachronic acid synthesis necessary for suppressing dauer formation, then the removal of the cholesterol precursor should not exacerbate the severity of the Daf-c phenotype in the presence of food. In both *C. elegans* and *P. pacificus*, 5 µg/ml cholesterol (∼13 µM) is routinely added to NGM medium [75, 76], whereas the removal of this exogenous cholesterol for two generations leads to constitutive dauer formation in the presence of food [50]. Indeed, we observed that wild-type *P. pacificus* form 68% active DL at 25°C when cultivated on NGM medium lacking the supplementary cholesterol in the medium, compared to 0% DL formation when raised with cholesterol. In contrast, cholesterol removal from *tu391* and *csu60* cultures grown on non-cholesterol supplemented NGM did not alter the percentage of DL (SI Fig. 5). The lack of response to cholesterol removal suggests that *Ppa-hsd-2* and the wild-type gene product of *tu391* are responsible for most of the metabolism of cholesterol in the biosynthesis of dauer-suppressing steroid hormones.

Utilizing whole genome sequencing of *tu391* and mapping lines, as well as previous fine mapping data [8], we confirmed that the mutation responsible for the *dfc-1(tu391)* Daf-c phenotype is located on Chromosome I, which contains 17 point mutations predicted to affect coding regions or splicing. To determine if the mutated coding sequence is represented in the transcripts, we amplified and sequenced mixed-stage *tu391* cDNA for a few candidates, including the patched-related protein PPA21795 (*Ppa-ptr-4*), and confirmed the point mutation in the transcript predicted to result in an amino acid substitution (Met>Lys). PPA21795 transcripts are also expressed at significantly lower levels in DL than in J2 and J3 larvae [74]. However, given the complexity of the PPA21795 exon-intron structure (33 exons), we did not pursue transgenic rescue experiments to test if it is responsible for the *tu391* phenotypes.

### *Ppa-hsd-2* influences amphid neuron identity

Given the finding that *P. pacificus* HSD-2 is upstream of the putative branching point for dauer formation and olfaction, we sought to investigate the possible neuronal changes associated with the enhanced attraction to ZTDO in *csu60* Daf-c adults. Using their connectivity profiles and axon trajectories, as well as cell-specific molecular markers, recent comparisons of the neuroanatomy between *C. elegans* and *P. pacificus* identified homologs of the two pairs of amphid olfactory neurons responsible for odor attraction, AWA and AWC [60]. The *P. pacificus* AWC homolog, AM7, expresses *Ppa-odr-7p::rfp* (versus the G-protein-coding *odr-3* in *C. elegans*), whereas the AWA homolog, AM3, expresses *Ppa-odr-3p::rfp* (versus the nuclear hormone receptor-coding *odr-7* in *C. elegans*). We found that while *Ppa-odr-7p::rfp* expression is unaltered in *csu60* mutants (SI Fig. 6), *Ppa-odr-3p::rfp* expression is dramatically altered in *csu60* in all post-hatching stages (Fig. 7). Specifically, in addition to the robust *Ppa-odr-3p::rfp* expression in AM3(AWA) and the weaker expression in AM4(ASK), 68% of *csu60* J4 larvae and adults also showed unexpected expression in a third pair of amphid neurons posterior to the AM3(AWA) (Table 2; Fig. 7A-B). The level of expression for each neuronal pair was also stronger in *csu60* neurons compared to wild type. It is not clear if the expression represents a duplication of the AM3(AWA) or the AM4(ASK) cell fate. To confirm that the increased *Ppa-odr-3* promoter activity and ectopic cellular expression correspond to increased *Ppa-odr-3* mRNA transcripts in *csu60*, we performed qPCR on mix-stage cultures and found that *Ppa-odr-3* transcripts are indeed ∼10 fold higher in *csu60* than in wildtype (SI Fig. 7).

**Figure 7.**
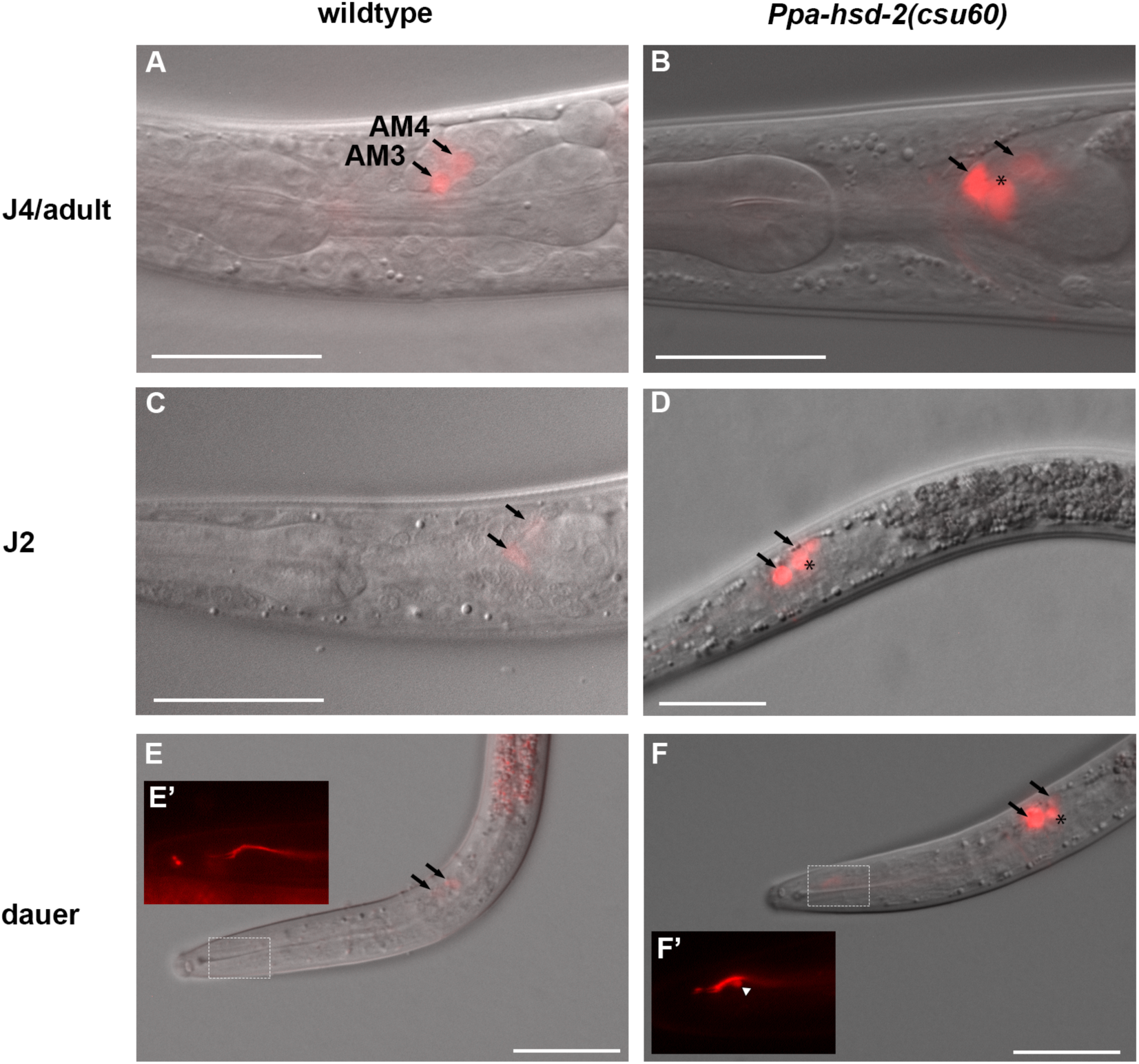
Ectopic expression of the *Ppa-odr-3p::rfp* reporter in amphid neurons in *Ppa-hsd-2(csu60)*. **(A-B)** Representative DIC and fluorescent overlaid images of J4 and adults. In wildtype, *Ppa-odr-3p::rfp* expression is found in AM3(AWA) and AM4(ASK) amphid neurons (arrows). In *csu60* mutants, reporter expression in AM3(AWA) and AM4(ASK) is stronger than wildtype and is found in an additional pair of amphid neurons posterior to the AM3(AWA)(*). **(C-D)** Ectopic *Ppa-odr-3p::rfp* expression is also found in *csu60* J2 larvae. **(E-F)** *Ppa-odr-3p::rfp* expression is also found in *csu60* DL. **(E’)** RFP fluorescence shows dual ciliated dendritic ending outlined in the dotted rectangular area of a different DL. **(F’)** RFP fluorescence shows dual ciliated dendritic ending outlined in the dotted rectangular area of the same DL. Triangle points to a type of posterior protrusion observed in 3 out of 17 *csu60* DL but not in wild-type DL (additional examples are shown in SI Fig. 8). Images of wild-type samples were taken at 700 ms exposure, and *csu60* samples were taken at 700 ms exposure or lower due to much stronger expression level. Sample sizes are shown in Table 2. Anterior is left and dorsal is up. Scale bar: 25 µm.

**Table 2.** Ppa-odr-3p::rfp expression in *Ppa-hsd-2(csu60)*

| Strain | Stage | No. Expression two pairs<br>AM3(AWA) + AM4(ASK) | No. Expression three pairs<br>AM3(AWA) + AM4(ASK) +<br>unknown (%) | N |
| --- | --- | --- | --- | --- |
| PS312 wildtype | J2 | 10 | 0 (0%) | 10 |
|  | Dauer | 22 | 0 (0%) | 22 |
|  | J4/adult* | 17 | 0 (0%) | 17 |
| *Ppa-hsd-2(csu60)* | J2** | 17 | 12 (41%) | 29 |
|  | Dauer | 24 | 4 (14%) | 28 |
|  | J4/adult | 10 | 21 (68%) | 31 |

\*An additional 22 adults with the same expression pattern were observed in Hong *et al*, *eLife*, 2019
\*\*Including some partial dauers that had darkened intestines but did not have buccal caps and radial constriction.
N: sample size

**Table 3:**
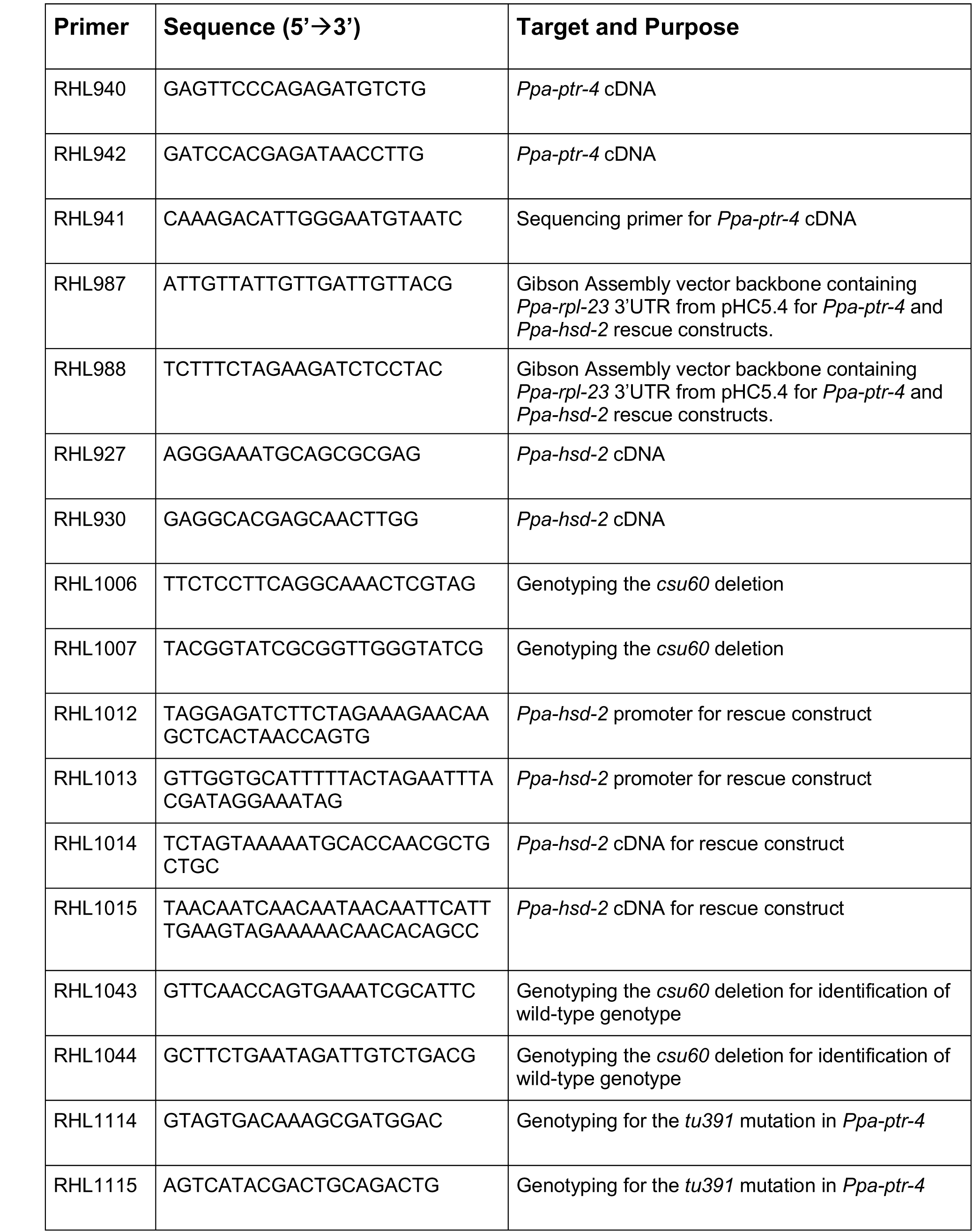
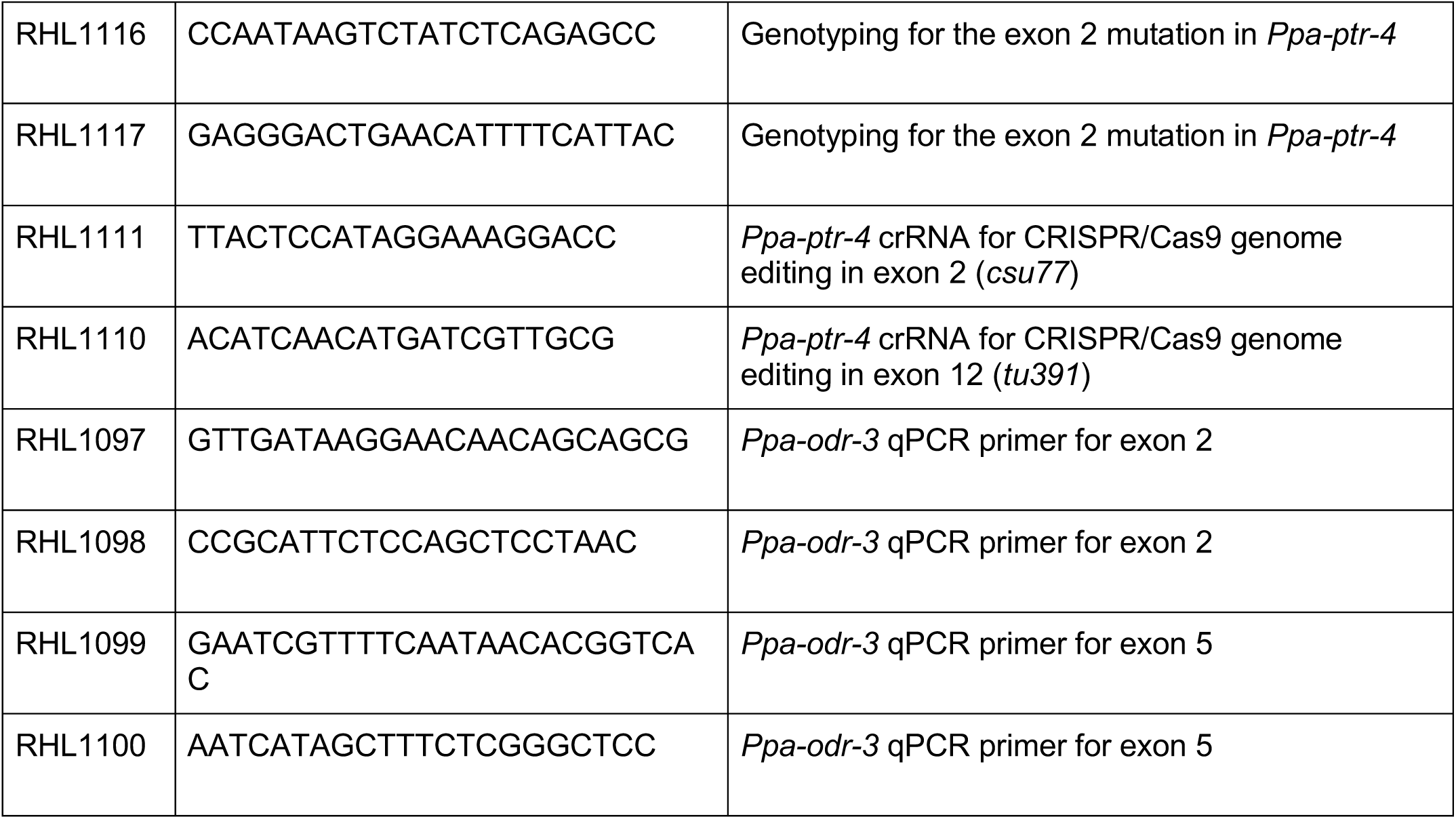
Primers

To determine if the duplicated *Ppa-odr-3p::rfp* expression approximated the frequency of Daf-c DL at 20°C, we also examined earlier developmental stages. In *P. pacificus*, the J1 larva undergoes a pre-hatching larval molt inside the eggshell, so the first active post-hatching stage is the J2 [77, 78]. Surprisingly, ∼41% of J2 larvae and ∼14% DL in *csu60* mutants, but 0% in wild type, also expressed ectopic *Ppa-odr-3p::rfp* in a duplicated pair of amphid neurons (Table 2) (Fig. 7C-F). This suggests that HSD activity is required to suppress a default-on *Ppa-odr-3-* expressing cell fate prior to hatching in the J1 larvae or during embryogenesis. The lower frequency of ectopic *Ppa-odr-3p::rfp* expression compared to the frequency of improper dauer entry decision for *csu60* may reflect the chimeric distribution of the reporter gene as an extrachromosomal array during development. Due to the higher RFP expression in *csu60* dauers, we observed the dual ciliated dendritic ends of the AM3(AWA) neurons, which was not visible in wild-type dauers [60]. These dendritic ends in *csu60* dauers also appeared to branch posteriorly (3 out of 17), which was not observed in the previous EM study on amphid dendrite morphology (Fig. 7E’-F’ and SI Fig. 8) [60]. However, it is unclear whether these posterior protrusions represent novel mutant dauer-stage structures or just rare wild-type variants previously not visible due to lower *odr-3p::rfp* expression in wild-type AM3(AWA). Because the duplicated *Ppa-odr-3p::rfp* pair was not found in wild-type DL, which have enhanced attraction to ZTDO, the difference in the *odr-3p::rfp* expression pattern alone cannot not fully explain the stronger ZTDO attraction in DL compared to adults in wild type, but is consistent with the possibility that the transformed wild-type amphid neuron may be involved in restricting dauer entry.

**Figure 8.**
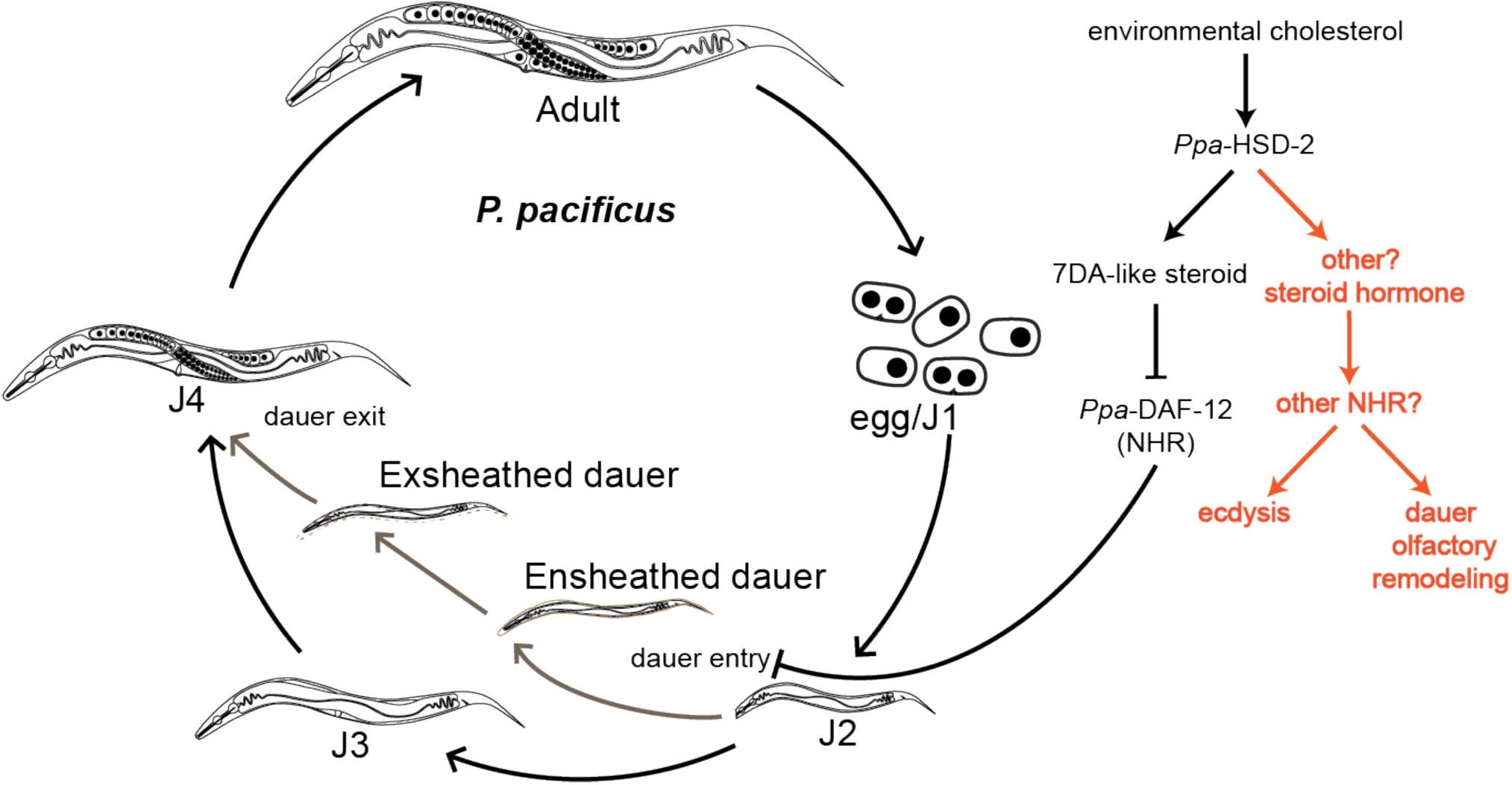
A model for the multiple roles of *Ppa-*HSD-2 in coordinating dauer entry, molting, and olfaction. From the cholesterol precursor, *Ppa-*HSD-2 is required for the synthesis of a Δ7-dafachronic acid-like steroid hormone (7DA). Binding of Δ7-DA to the DAF-12 nuclear hormone receptor represses decision for dauer entry. *Ppa-*HSD-2 may also be required for producing other steroid hormones that can explain the DAF-12-independent phenotypes of defective ecdysis during reproductive development and enhanced host odor attraction in the Daf-c adults.

## Discussion

In order to effectively compare the olfactory behaviors of adults versus host-seeking dauer larvae, we have characterized the first two Daf-c alleles in the entomophilic nematode *P. pacificus*. The *csu60* Daf-c phenotypes are due to a deletion in the sole HydroxySteroid Dehydrogenase (HSD) in *P. pacificus*. This gene is likely responsible for the synthesis of a Δ7-DA-like steroid ligand, which regulates the conserved nuclear hormone receptor DAF-12 (Fig. 8). In *C. elegans*, orthologous mutations in the HSD homologs on their own do not cause constitutive dauer or molting defects, which suggests that steroid hormone signaling in *P. pacificus* represents an ancestral state more in common with parasitic nematodes than with *C. elegans*. Most notably, the complete loss of *Ppa-hsd-2* resulted in host pheromone attraction by adults in a *daf-12*-independent manner, and is accompanied by early neuronal fate changes that lead to ectopic *Ppa-odr-3* G-protein expression in a pair of chemosensory amphid neurons. By testing two well-studied Daf-c mutants in *C. elegans*, we found that Daf-c alleles lowered the already reduced wild-type DL response to adult attractants. This is in stark contrast to the *P. pacificus* Daf-c alleles, which did not affect the response of DL but instead increased the attraction of adults to host odorants. Thus, regulators of dauer formation affect the olfactory system in developmentally distinct manners for divergent species of nematodes.

### Dafachronic acids control cuticle exsheathment and dauer entry

Retention of the previous cuticle, or ensheathment, is a hallmark of the infective larvae of many parasitic nematodes as well as insect larvae. In insects, the pharate larva forms a new cuticle while still within, but separated from, the old cuticle of the previous stage [79]. In nematodes, the exsheathment of infective larvae in mammalian parasites and entomopathogenic nematodes (EPN) often marks the transition to the parasitic phase of the life cycle. Cues for exsheathment are a combination of intrinsic physiological factors, such as the time since emergence from the host, and extrinsic environmental factors, such as temperature or host cues [53,80–82]. For example, in the IJs of the EPN *Heterorhabditis bacteriophora*, exsheathing the previously attached old J2 cuticle occurs upon entering an insect host, while in another EPN, *Steinernema carpocapsae,* IJs are unresponsive to host cues for exsheathment [53]. In the pathogenic nematode of sheep, *Haemonchus contortus*, exogenous application of an endogenous Δ7-DA promotes the exsheathment of infective L3 larvae [10]. Although there are no candidate genes known to control dauer exsheathment in *C. elegans–* because its dauers are ‘naked’*–* conserved genes involved in molting may nevertheless modulate the timing of cuticle shedding. Only one uncloned Daf-c allele, *daf-13(m66)*, has been reported to display a dauer-specific cuticle defect [83]. However, we could not observe either the Daf-c phenotype or the cuticle defect from a revived stock. In all four *C. elegans* molts, the Zona-Pellucida domain protein FBN-1 and integrins make up a part of the sheath between the old and new cuticles [84]. We speculate that a similar mechanism may be involved in *P. pacificus* such that the early detachment of the old cuticle in the Daf-c DL leads to loss of contact with the mouth plug (and other orifices), resulting in the incarceration of the DL inside the old J2 cuticle. Because dauer entry involves the formation of a buccal cap, the retention of the J2 cuticle in *P. pacificus* DL may render Daf-c DL especially prone to the incarcerated exsheathment phenotype by disrupting the timing of these highly choreographed events.

### Steroid hormones directly regulate neuronal cell fate

The ectopic expression of *Ppa-odr-3p::rfp* in *csu60* J2 animals shows that steroid hormones in *P. pacificus* are also responsible for maintaining proper cell fate in putative olfactory neurons at or prior to the decision for dauer entry in J2 animals. The homeotic transformation of the AWB to AWC neurons due to mutations in the LIM-4 transcription factor was accompanied by changes in dendritic morphology and chemotaxis behavior [85]. Based on a similar rationale, the loss of *Ppa-hsd-2* could transform AM10(ASI) or AM6(ASG), which are normally involved in inhibiting dauer entry in *C. elegans*, into AM3(AWA) olfactory neurons. Coincidentally, AWA and ASG are sister cells in the *C. elegans* cell lineage (www.wormatlas.org). However, we did not observe fewer *Ppa-che-1p::rfp* expression in the amphid neurons of *Ppa-hsd-2(csu60)* animals, as would be expected if one pair of the neuron mis-expressing *Ppa-odr-3p::rfp* was derived from the AM6(ASG) or AM5(ASE) neurons [60] (SI Fig. 6). Given the dynamic transcriptional profiles associated with the dauer stage in *P. pacificus* [86], Daf-c mutants have the potential to identify unknown molecular pathways involved in dauer commitment outside of *C. elegans*.

### Tributaries of DAF-12 flow into differing gene regulatory networks

Microarray and RNA-seq studies have shown substantial divergence of dauer-specific genes between *P. pacificus* and *C. elegans*, as well as more upregulated dauer-specific genes in *P. pacificus* than in *C. elegans* [74, 86]. In *P. pacificus*, the DAF-12/Δ7-DA module is a shared regulatory target for both dauer entry and mouth form dimorphism [37, 87]. The dye-filling defect in *tu391* J2 and adults, but not in the DL, suggests that defects in the amphid neurons, possibly those responsible for negatively regulating dauer entry, are remodeled during the dauer stage. The *C. elegans daf-19* gene, which encodes the RFX family of transcription factors, is required for sensory neuron cilium formation and is one of the few mutations that results in both Daf-c as well as dye-filling defective phenotypes [62–64]. Although *dfc-1(tu391)* shares the Daf-c and dye-filling defective (in the non-dauers), *dfc-1(tu391)* differs from *C. elegans daf-19* mutants in that they do not exhibit amphid sheath defects [88] and lack a chemosensory defect. Moreover, mutations in the *P. pacificus daf-19* ortholog did not result in a dauer formation defect but did show a dye-filling phenotype [47]. The particular combination of phenotypes in *dfc-1(tu391)* has no obvious candidates.

In *C. elegans*, genetic dissection of the relative contribution of the steroidogenic enzymes is confounded by the unknown roles the HSD-2 and HSD-3 paralogs play in dafachronic acid biosynthesis. HSD-1 is a component in one of the two branched pathways that includes the cytochrome P450 DAF-9 required for dafachronic acid biosynthesis [89]. While HSD-1 and DAF-9 use cholesterol precursors to synthesize Δ4-DA, the Rieske-like oxygenases DAF-36 and DAF-9 synthesize Δ7-DA, and together the dafachronic acid hormones converge to promote reproductive development by binding to the DAF-12 receptor in the inactive state [15,73,89]. Mutations in *hsd-1* contribute to a nearly fully penetrant Daf-c synthetic phenotype only when combined with a second mutation in NCR-1/cholesterol transporter, DAF-36/Rieske-like oxygenase, or DAF-28/insulin [73]. In *P. pacificus*, the nearly fully penetrant constitutive dauer phenotype of the *csu60* null allele at 25°C suggests that the absence of *Ppa-hsd-*2, rather than a temperature sensitivity of its gene product, can lead to an almost complete de-repression of dauer formation in a *daf-12*-dependent manner. Because the DNA lesion for *tu391* is not yet known, it is difficult to speculate if the *tu391* mutation acts in the same pathway as *Ppa-hsd-2*. With only a single *hsd* in the *P. pacificus* genome, future studies will likely reveal the degree of functional overlap the *P. pacificus ncr-1, daf-36,* and *daf-9* orthologs have with *Ppa-hsd-2* in dauer regulation and olfactory behavior.

### *C. elegans* dauers have attenuated responses to adult odors

To achieve organismal plasticity, it is has been shown that *C. elegans* dauer entry involves the remodeling of glia and neural circuitry, as well as differential expression of chemosensory and neuropeptide receptors [3,4,6,7,19]. While dauer-induced remodeling of the AWC dendritic ends that increase their surface area could heighten sensitivity, our results show that wild-type, *daf-2,* and *daf-7* DL were actually less attracted than adults to several odors, and the DL of two *daf-7* alleles even displayed avoidance of isoamyl alcohol. Mutation of the *daf-2* gene also mostly eliminated acute CO_2_ avoidance [20], suggesting that Daf-c mutations have multiple effects on chemosensation. Other dauer-specific changes in amphid gene expression and function have been observed. Two pairs of amphid neurons, the AWA and AWC, are the primary mediators of chemotaxis toward attractive odorants [24]. In *C. elegans* adults, the AWA neurons detect 2,3-butanedione (diacetyl), while the AWC neurons sense isoamyl alcohol and 2,3-pentanedione. A significant upregulation of neuropeptide expression during dauer entry occurs in both *C. elegans* and *P. pacificus* [86], some of which may correlate with stage-specific olfactory behavior, as well as dauer-specific behaviors such as nictation and the change in CO_2_ response valence [6]. A combination of cellular identity and gene expression changes may be responsible for stage-specific chemosensory behaviors between adults and DL [69, 70].

Our study shows that dauers in two divergent nematodes can have either attenuated or enhanced odor responses relative to reproductive adults, and that the differences can be intensified in the Daf-c mutants. However, the genetic mechanisms coordinating these developmental and behavioral changes in these two species appear to evolve in separate gene regulatory networks. Several recent *C. elegans* studies on the remodeling that occur after entry into dauer and in post-dauers adults show significant changes to olfaction and foraging behavior [19,69,70]. Our study also revealed that *Ppa-hsd-2(csu60)*; *daf-12(tu389)* mutants have enhanced adult odor attraction, along with a mild but noticeable ecdysis defect in non-dauer larval stages. This finding invigorates the suspicion that *Ppa*-HSD-2 may also be involved in the production of other steroid hormones, such as ligands targeting the two putative *P. pacificus* ecdysone receptors [90]. Although both *Caenorhabditis* and *Pristionchus* species co-occur on rotten vegetation as both DL and feeding stages [91], only DL from *Pristionchus* species have been isolated from beetles [32, 34]. The robust chemoattraction to host pheromone shown in this study supports the model that the DL is the preferred host-seeking stage capable of locating beetle hosts. We expect we will have a better understanding of how the decision for dauer entry is genetically coordinated with host-associated remodeling of behavior when the cognate olfactory neurons for the host pheromones are identified in *P. pacificus*.

## Materials and Methods

### Nematode Strains

*P. pacificus* and *C. elegans* strains were maintained at 20°C on NGM plates seeded with OP50 as described previously [68]. The following *P. pacificus* strains were used: PS312 (California wild-type, synonymous with RS2333), *Ppa-obi-1(tu404)* ChrI, *Ppa-dfc-1(tu391)* ChrI, and *csu60* ChrII; *Ppa-daf-12(tu389)* ChrX. For *C. elegans*, wild-type N2, as well as Daf-c mutants *daf-2(e1370)* and *daf-7(ok3125, e1372)* were used. A previous screen for Daf-c mutants using 50 mM ENU or EMS yielded two highly penetrant alleles *dfc-1(tu391)* and *Ppa-dfc-2(tu392)*, but only *dfc-1(tu391)* was viable for further characterization [8], while *csu60* was isolated as an off-target mutant from a CRISPR/Cas9-mediated mutagenesis against a TGF-ß ortholog (Contig115-snapTAU.31)). Each allele was backcrossed three times to PS312 for the Daf-c phenotype before further characterization. F_1_ complementation cross between *dfc-1(tu391); Ppa-daf-6p::rfp* and *csu60* did not result in RFP+ F_1_ DL on food. Double mutants with *obi-1(tu404)* were initially selected for the long body phenotype of *obi-1* along with the Daf-c phenotype to create *rlh177* (*tu391*; *tu404*) and *rlh192* (*csu60*; *tu404*). Each double mutant with *obi-1(tu404)* was confirmed by sequencing the single nucleotide deletion in *tu404.* The cloning and construction of the *Ppa-daf-6p::rfp* (*tuEx231*), *Ppa-odr-3p::rfp* (*tuEx265*), *Ppa-odr-7p::rfp*(*tuEx297*), and *Ppa-che-1p::rfp* (*lucEx367*) transgenic lines were described elsewhere [60]. To examine the expression of these amphid sheath and neuronal markers in the *csu60* background, we crossed RFP-expressing transgenic males into *csu60* J4 hermaphrodites, and then established individual transgenic lines from RFP+ F_2_ progeny with the Daf-c phenotype.

For the epistasis tests, we first crossed *Ppa-daf-12(tu389)* hermaphrodites with *csu60/+* males, then isolated F_2_ progeny homozygous for the *csu60* deletion by PCR genotyping and for the single nucleotide substitution of *Ppa-daf-12(tu389)* by Sanger sequencing [8]. The double mutant (*csu60; tu389*) did not produce DL on well-fed or on starved plates, indicating the Daf-d phenotype of *Ppa-daf-12(tu389)* is epistatic to the Daf-c phenotype of *csu60*. Because the genetic lesion responsible for *tu391* was less obvious (i.e. not a large deletion), we examined the segregation pattern of Daf phenotypes in addition to the genotype of the *Ppa-ptr-4* candidate locus. We constructed a double mutant strain heterozygous for the *daf-12(tu389)* nonsense mutation (checked by sequencing) that segregated for the Daf-c phenotype in the F_2_ progeny, which could be either heterozygous or homozygous for the *tu391* allele. We then examined the F_3_ progeny for dauer formation defects guided by the following rational: If the *daf-12* Daf-d phenotype is epistatic to *tu391,* then no progeny of a non-DL F_2_ homozygous for *daf-12(tu389)* will become DL. Alternatively, if the *tu391* Daf-c phenotype is epistatic to *daf-12,* then some F_3_ Daf-c progeny will be homozygous for *daf-12(tu389)*. We found that 0 out of the 5 F_2_ lines from Daf-c dauers that exited were homozygous for the *Ppa-daf-12(tu389)* genotype (χ^2^, P<0.0001), and none of the 6 F_2_ lines homozygous for *Ppa-daf-12(tu389)* segregated for constitutive DL (χ^2^, P<0.0001). Furthermore, all 6 of these *daf-12(tu389)* lines were Daf-d as well as homozygous for the *Ppa-ptr-4* mutation, suggesting that *daf-12* also acts downstream of *Ppa-dfc-1(tu391)* for dauer regulation.

### Temperature and cholesterol sensitivity

For each experiment, 10 gravid hermaphrodites were picked onto freshly seeded 6 cm NGM plates and incubated at 15°, 20°, or 25°C. Plates were scored for third-stage larvae larvae (active dauer, incarcerated dauer, J3 larvae) after 5 days for Daf-c or 3 days for wildtype PS312 (because they starve after 5 days under the same condition). To determine how long the ensheathed DL remain incarcerated, we transferred the immobilized DL at 20°C onto new OP50 NGM plates. After one day at 20° or 25°C, these cultures were scored for the presence of active DL and incarcerated DL. To determine response to the removal of cholesterol, which is usually supplemented in the NGM (1 µg/ml), we cultivated worms on NGM lacking cholesterol for three generations in the same manner before scoring for DL as described above for temperature sensitivity.

### Exogenous Δ7-dafachronic acid

10 µl of 75 µM Δ7-dafachronic acid or 100% ethanol vehicle control was mixed into 90 µl OP50 before seeding onto NGM plates. After two days, we picked 15-20 *tu391* or *csu60* gravid hermaphrodites onto each plate, which were allowed to lay eggs for 5 hours before being removed for egg synchronization, or if the hermaphrodites lay fewer than 50 eggs they were instead removed after 24 hours. Plates were incubated at 20°C or 25°C for 4-5 days and then scored for dauer larvae and non-dauer stages.

### Behavioral assays

Pharyngeal pumping rates were observed through a Leica DM6000 upright microscope with a 40x objective, without any anesthetic. J3 larvae, dauer larvae, and incarcerated dauer larvae were mounted onto M9 buffer and 2% agar pads on microscope slides. We counted at least four 15-second intervals per animal, which were then summed to obtain pumps/minute.

The population chemotaxis assay for *P. pacificus* were performed on covered 10 cm Petri dishes for ∼16 hours at 22°C as previously described [39, 68]. However, because *P. pacificus* DL is coated with a long-chain polyunsaturated wax ester (nematoil) that makes them highly hydrophobic [35], we modified the chemotaxis assay by adopting a chemotaxis medium containing a detergent (10 mM MOPS pH 7; 2.5% Tween 20; 1.5% Bacto-agar) [65]. This MOPS/Tween medium promotes both adult and DL dispersal, which when 5% rather than 10% ZTDO was used as an attractant, likely contributed to the slightly higher CI values for PS312 and *obi-1(tu404)* adults than expected when compared to previously reported [68]. DL were loaded to the center of the plate by placing and removing immediately a 1 cm^2^ agar chunk from a culturing plate containing either wild-type dauers from starved cultures or Daf-c DL from densely populated cultures. Young adult worms from nearly-saturated cultures were washed twice with M9 buffer and collected by centrifuging at 2500 rpm for 2 minutes. Approximately 100 worms were loaded onto each assay plate. Assays with *obi-1(tu404)* DL could not be successfully performed due to their fragility and higher mortality. To anesthetize animals at the odor sources, we place 1.5 µl of 1 M sodium azide on both sources and used 100% ethanol as the counter attractant. All odorants were diluted with ethanol to the following concentrations: 5% (*Z)*-7-tetradece-2-one (previous studies used 10% ZTDO, which resulted in more DL paralysis at loading), 1% methyl myristate, 1% (*E)*-11-tetradecenyl acetate, 1% (*E)*-11-hexadecenal, 10% ß-caryophyllene, 1% 2,3-butadione(diacetyl), 1% 2,3-pentadione, and 1% isoamyl alcohol. Interestingly, we found sodium azide to be superfluous when using ZTDO as the attractant due to ZTDO’s paralyzing effects on DL. Chemicals were purchased from Fisher Scientific, Sigma-Aldrich, or Bedoukian Research (Danbury, CT).

The chemotaxis assays for *C. elegans* were set up in a quadrant with worms loaded onto the center of 6 cm plates containing the same MOPS/Tween agar as described above, along with two opposing pairs of odors and controls [92, 93]. 0.5 µl of 1 M sodium azide was spotted onto each odor or counter-attractant. Since *C. elegans* DL are hydrophilic, we collected DL primarily from the condensations present on the underside of the lids of recently starved plates by washing the lids with sterile spring water (Arrowhead, CA). For assays with Daf-c mutants, each plate was scored for either DL or YA using visual confirmation (each plate can only be scored for one stage). Chemotaxis assays lasted 1-1.5 hours.

### CO_2_ Chemotaxis assays

Young adults were washed off plates using M9 buffer and collected in a 65 mm-Syracuse watch glass. Animals were then washed 2X with M9 buffer followed by 1X with sterile water and immediately transferred to the center of 6 cm MOPS/Tween agar plates. Excess water was removed with Whatman paper. DL were directly transferred to the 6 cm MOPS/Tween agar plates on chunks of agar from starved plates. A CO_2_ gradient was generated by delivering specific compositions of gasses through holes in the plate lids as previously described [94]. A certified gas mixture containing 21% O_2_, balance N_2_ (Airgas) air control was pumped through one hole whereas a certified gas mixture containing 10% CO_2_, 21% O_2_, balance N_2_ (Airgas) was pumped through the other hole. Gasses were pumped through ¼-inch flexible PVC tubing using a syringe pump (PHD2000, Harvard Apparatus) at a flow rate of 0.5 ml/min. Adults and DL were allowed to migrate in the CO_2_ gradient for 1 hour and 90 minutes, respectively. Scoring regions were defined as opposite halves of the assay plate 1cm away from the center line (as shown in the schematic). At the end of the assay, animals that migrated to opposite scoring regions were scored according to the formula:

Chemotaxis index (CI) = [(# animals at CO_2_ side) – (# animals at air side)] / (Total # animals at both sides).

As a control for directional bias, two assays were always run simultaneously with CO_2_ gradients in opposite directions. Assays were discarded if the difference in the CI between the two plates was ≥0.9 or if fewer than 7 animals moved to both scoring regions combined on either assay plate.

### DiI staining

Bacteria were removed from worm cultures by washing with M9 buffer followed by centrifuging. The cleaned worms were soaked in 300 µl 6.7 mM Vybrant™ DiI Cell-Labeling Solution (Molecular Probes V22889) diluted in M9 with gentle inversion every 30 minutes away from direct light. After 3 hours, worms were washed twice with 5x volume M9, followed by de-staining for at least one hour on plates seeded with OP50. To isolate and stain DL, 0.1% SDS was added to the M9 used in all steps. DiI staining was observed using a Leica DM6000 with a fluorescent microscope. While six pairs of the amphid neurons have been observed to take up DiI in wild-type PS312 [60], we scored for dye uptake in the mutants only in the most robustly stained pairs– AM1, AM2, AM4 – that correspond to the *C. elegans* neuronal homologs ASH, ADL, and ASK, respectively.

### *Ppa-hsd-2* transgenic rescue of *csu60*

15-20 gravid *csu60* hermaphrodites with or without the *Ppa-hsd-2* transgenic rescue construct were picked onto plates with food, and allowed to lay eggs for 24 hours before removal to obtain a roughly synchronized population of embryos. The plates were incubated at 20°C for 5 days, then scored for dauer larvae and non-dauer stages, as well as for the expression of the co-injection marker *Ppa-egl-20p::rfp* as a proxy for the presence of the *Ppa-hsd-2* rescue transgene.

Interestingly, we found that this method of synchronizing embryos led to a higher percentage of the population remaining as DL, compared to not synchronizing the embryos used in the temperature sensitivity assay.

### Molecular Cloning

To identify the mutations, we first rough mapped the mutation to a chromosomal region using Simple Sequence Length Polymorphic markers (SSLP) on 45 mapping lines generated with the mapping strain RS5278 (Bolivia). Each mapping line was a descendent of a single F_2_ dauer larva on non-starved F1 plates that exited when we transferred it to a fresh plate of OP50. Nematodes were washed individually from each mapping line in approximate equal pellet sizes prior to pooling and genomic DNA extraction using the MasterPure Kit (Epicentre). We sequenced the whole genomes of *tu391* and *csu60*, as well as additional mapping lines for fine mapping of *tu391* and *csu60*, respectively [95] (∼73 million 100 nt reads for each genome and bulk mapping samples; BGISEQ-500 platform with DNA Nanoball technology, BGI America). For *tu391*, our fine mapping results with 22 mapping lines concurred with the finding by Ogawa *et al* (2009) that the mutation lies on Chromosome I between S563 and S440, and identified 6 candidate point mutations within this ∼2.4 Mbp region. For *csu60,* rough mapping indicated linkage to Chromosome II, with the additional fine mapping of 47 recombinant lines and the *csu60* genome sequencing revealed a large 95 kb deletion on Chromosome II (1370000-1465000) that encompassed 11 open reading frames. One of the predicted genes in this interval, PPA10139 (UMM-S10-1.4-mRNA-1), encodes a 376 amino acid homolog of the *C. elegans* hydroxysteroid dehydrogenases 2 (HSD-2).

To make the *Ppa-hsd-2* cDNA rescue transgene, 2063 bp of the *Ppa-hsd-2* promoter was amplified from PS312 genomic DNA using RHL1012 and RHL1013. The full-length *Ppa-hsd-2* cDNA sequence was amplified from PS312 mixed stage cDNA using RHL1014 and RHL1015. The *Ppa-hsd-2* promoter and cDNA were combined with *Ppa-rpl-23* 3’ UTR to make pHC12. The mix injected into PS312 adult hermaphrodites contains the following: 2.5 ng/µl pHC12, 4 ng/µl *Ppa-egl-20p::rfp*, 80 ng/µl *csu60* gDNA (all digested with PstI). The transgenic line was then crossed into the *csu60* mutant background using PCR to genotype for the deletion (RHL1006 and RHL1007 will amplify 693 bp if there is a deletion; RHL1043 and RHL1044 will amplify 1008 bp if the same locus is wildtype) and scored as a percentage of dauer or non-dauers (J3/J4/adult) containing the rescue transgene. The *csuEx54* transgene transmission rate was approximately 27%.

### Reverse transcription and Quantitative Real-Time PCR (qPCR)

We used two pairs of amplicons with each primer spanning over exon boundaries to determine the relative level of *Ppa-odr-3* transcript expression in *csu60.* cDNA was synthesized using random hexamers (N6) and polyT primers using ∼500 ng total RNA from mix staged worms (Thermo Fisher Maxima, M1681). We performed two RNA extractions and three cDNA synthesis to run 12 technical replicates for each of the primer sets. We ran the qPCR reactions using 1-2 µl cDNA in SYBR Green Master Mix (Biorad) totaling 20 µl in a Biorad CFX96 machine using 60°C as the annealing temperature. *Ppa-*beta-tubulin and *Ppa-cdc-42* (RHO GTPase) were used as housekeeping genes and the relative fold change in expression between wildtype and *csu60* was calculated using the delta delta C_t_ method [96, 97].

### Phylogeny

The amino acid sequences of potential HSD homologs were first identified by BLASTX searches on WormBase. The phylogeny trees were built using the entire amino acid sequences by first making an alignment and removal of positions with gap using T-COFFEE, followed by Maximum Likelihood phylogeny by PhyML, and finally tree rendering by TreeDyn (www.phylogeny.fr)[98]. Midpoint rooting was used and branch support ≥50% is shown.

### Statistical analysis

Statistical tests were analyzed using GraphPad Prism 8 (La Jolla, CA) and Microsoft Excel. Values were checked for normal distribution by the Kolmogorov-Smirnov test before applying either multiple testing for parametric (Sidak, Tukey, or Dunnett) or non-parametric datasets (Kruskal-Wallis or Mann-Whitney).

## Acknowledgement

We thank C. Rödelsperger for whole genome sequence analysis, E. J. Corey for Δ7-DA by way of the Sommer lab, J. Ly and J. Cardenas for technical assistance, and the Developmental Biology class in Fall 2018 for *csu60* mapping. We also thank the *C. elegans* Genetic Center for the *daf-13(m66)* strain and T. Baiocchi for helpful comments on the manuscript. Funding is provided by NIH SC3GM105579 (to RLH), NIH 5T34GM008395-29 (to RV), NIH R01 441802-HE-31169 (to EAH), and NIH F32Al147617 (to NB).

## Abbreviations

DL: dauer larva

ZTDO: (*Z*)-7-tetradecen-2-one

## Supplementary Figures

**SI Figure 1.**
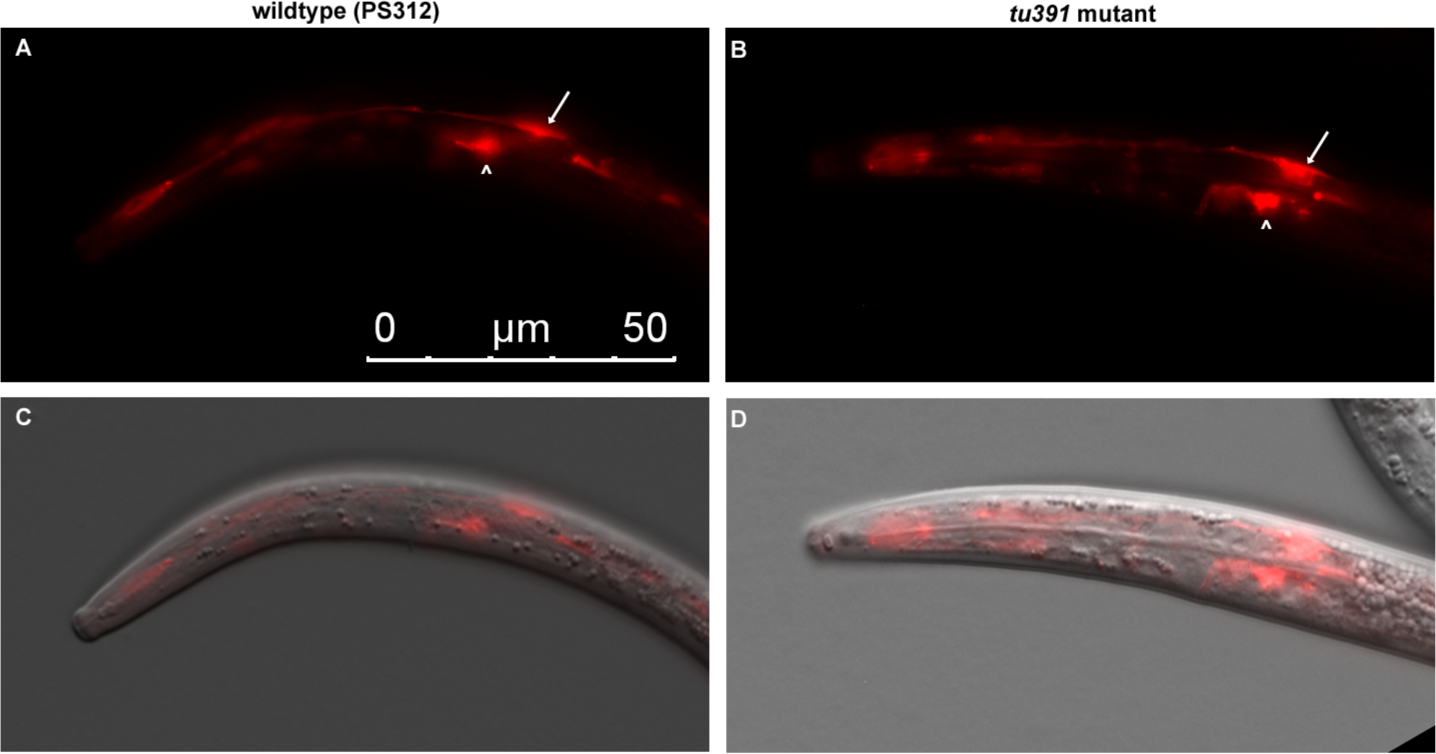
*tu391* dauer larvae have normal amphid sheath morphology. Fluorescent and DIC overlay images of wild-type PS312 **(A, C)** and *tu391* **(B, D)** dauers show similar expression profiles of the amphid sheath marker *Ppa-daf-6p::rfp* (RS2818) indicated with arrow heads. “^” marks the excretory cell. Anterior is left and dorsal is up. The scale bar in A represents all panels.

**SI Figure 2.**
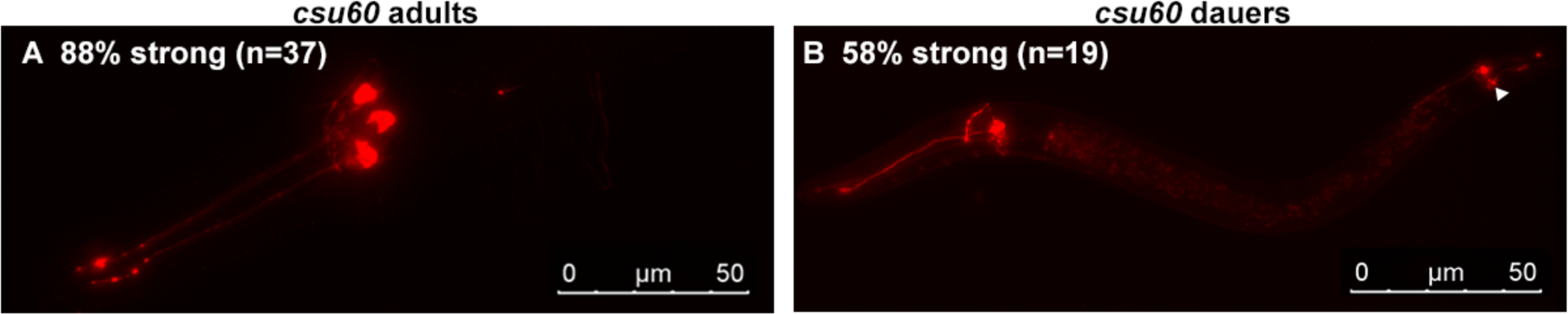
***csu60* is wild type in live dye DiI uptake**. **(A)** Adult and **(B)** dauer larvae take up dye at a level comparable to wildtype (Fig. 2). Max projections of stacked images, exposure 200 ms.

**SI Figure 3.**
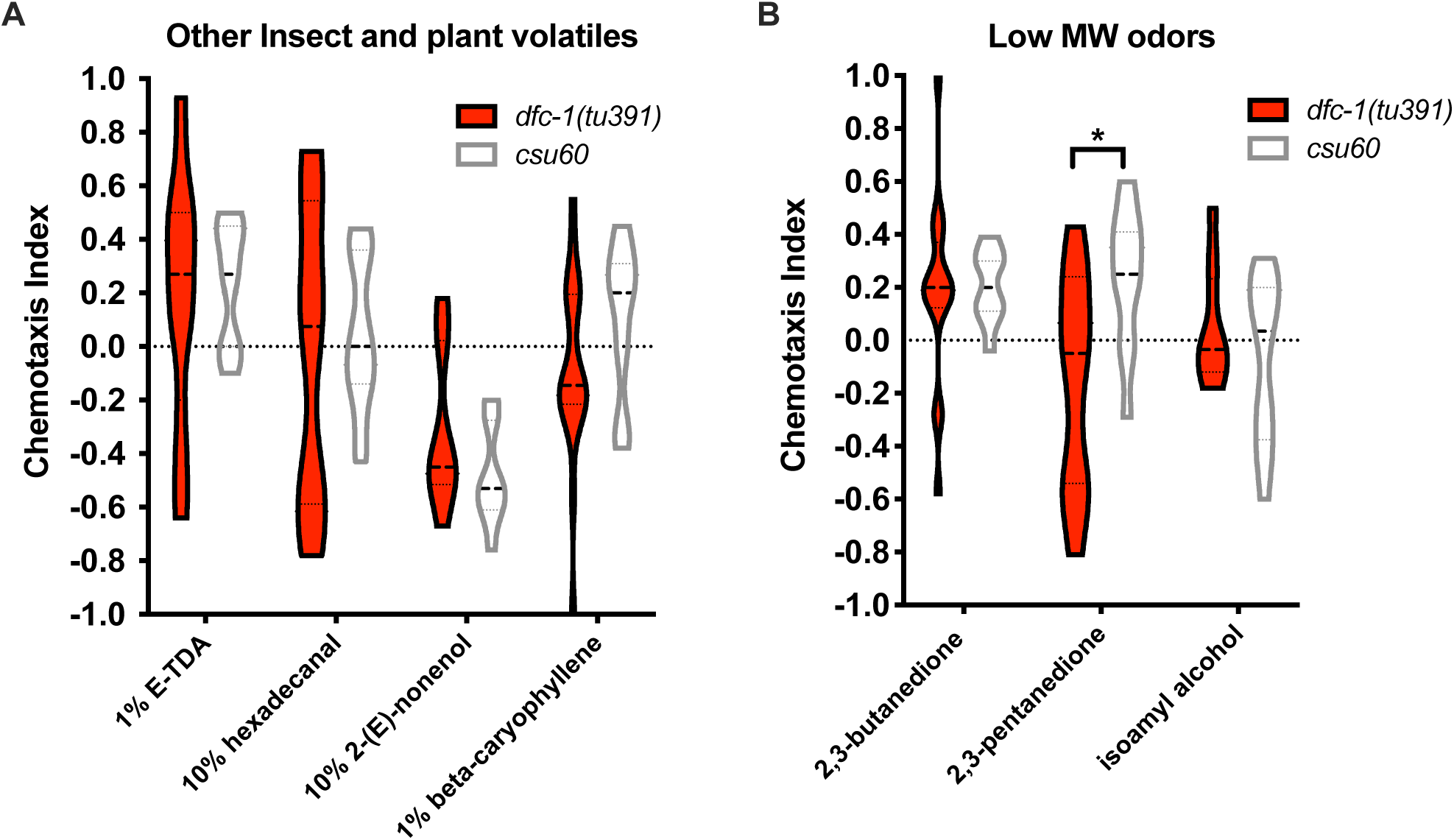
Olfactory responses of Daf-c dauer larvae. (A) Dauer larvae of Daf-c mutants do not respond to other insect pheromones known to be attractive to adults (1% *E*-11-tetradecenyl acetate (E-TDA), 10% hexadecanal, 10% 2-(*E*)-nonenol) and the plant defense volatile 1% ß-caryophyllene (Hong *et al*, *Current Biology*, 2006). **(B)** Dauer larvae of Daf-c mutants also do not respond to small molecular weight compounds that are strong attractants to *C. elegans*: 1% 2,3-butanedione (diacetyl), 1% 2,3-pentanedione, and 1% isoamyl alcohol. Two-way ANOVA with Sidak’s multiple comparisons test between *tu391* and *csu60*. Violin plots show medians with quartiles. \**P*<0.05.

**SI Figure 4.**
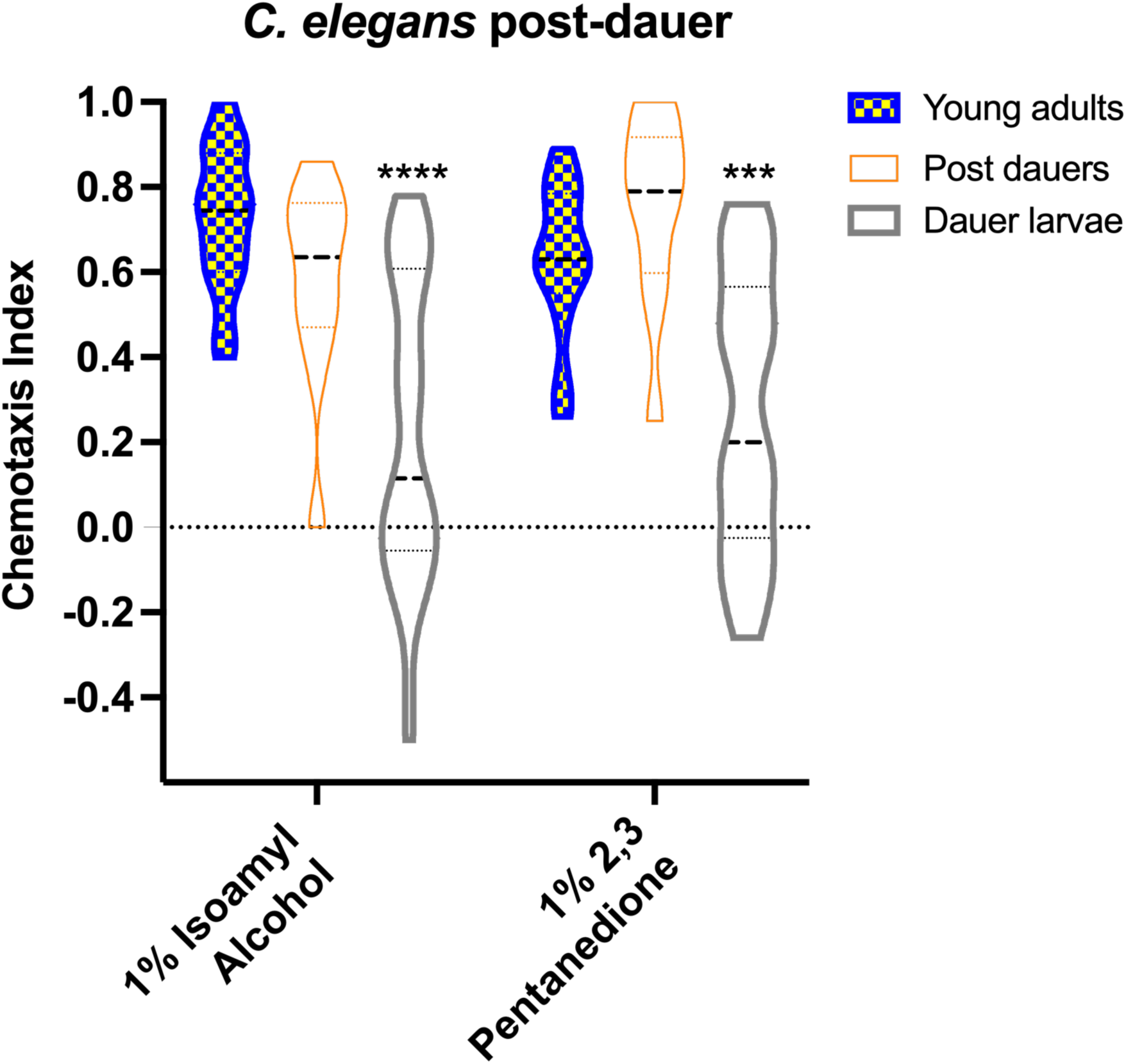
**Olfactory response of *C. elegans* post-dauer adults.** The chemotaxis response of wild-type post-dauer young adults in *C. elegans* resemble never-dauer young adults but not dauer larvae. IAA: One-way ANOVA against adults with Kruskal-Wallis test. Pentanedione: One-way ANOVA against adults with Sidak’s post-hoc test. \*\*\**P*<0.001; \*\*\*\**P*<0.0001.

**SI Figure 5.**
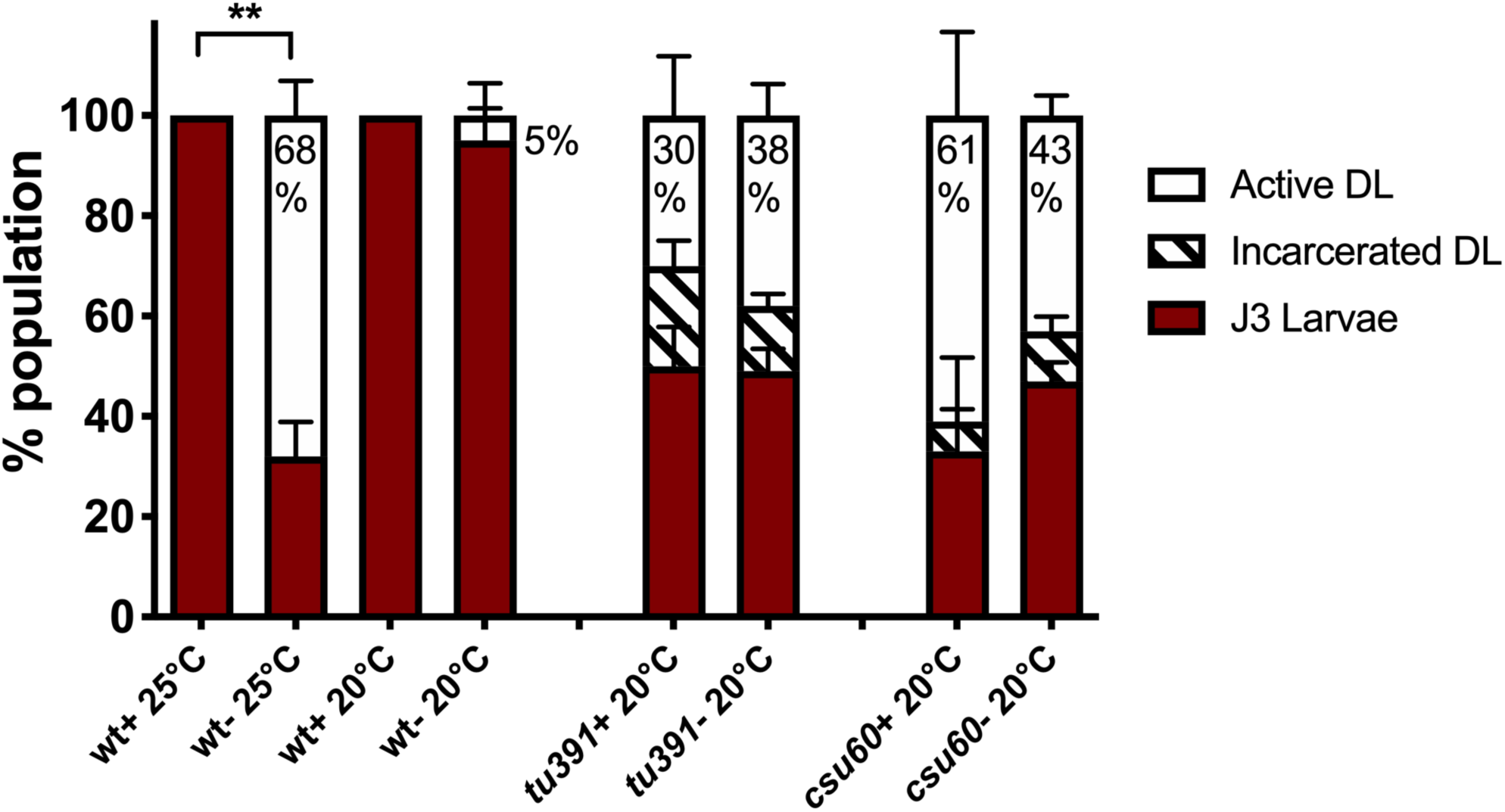
**Effects of cholesterol removal in Daf-c mutants.** The progeny of worms that were maintained for two generations on NGM plates with (+) or without (-) the normal cholesterol supplement in the medium were scored for the formation of active DL. At 25°C, the removal of cholesterol greatly increased dauer formation in wildtype worms but not in Daf-c mutants at 20°C (Comparisons were done at 20°C, since >92% of *tu391* and *csu60* form DL already at 25°C). The percentage of active DL is indicated. For each condition, 4-5 assays were performed over two trials. Difference in active DL in wildtype: One-way ANOVA with Kruskal-Wallis test. \*\**P*<0.01. Difference in active DL in each Daf-c mutant: t-test with Mann-Whitney test detected no significant difference.

**SI Figure 6.**
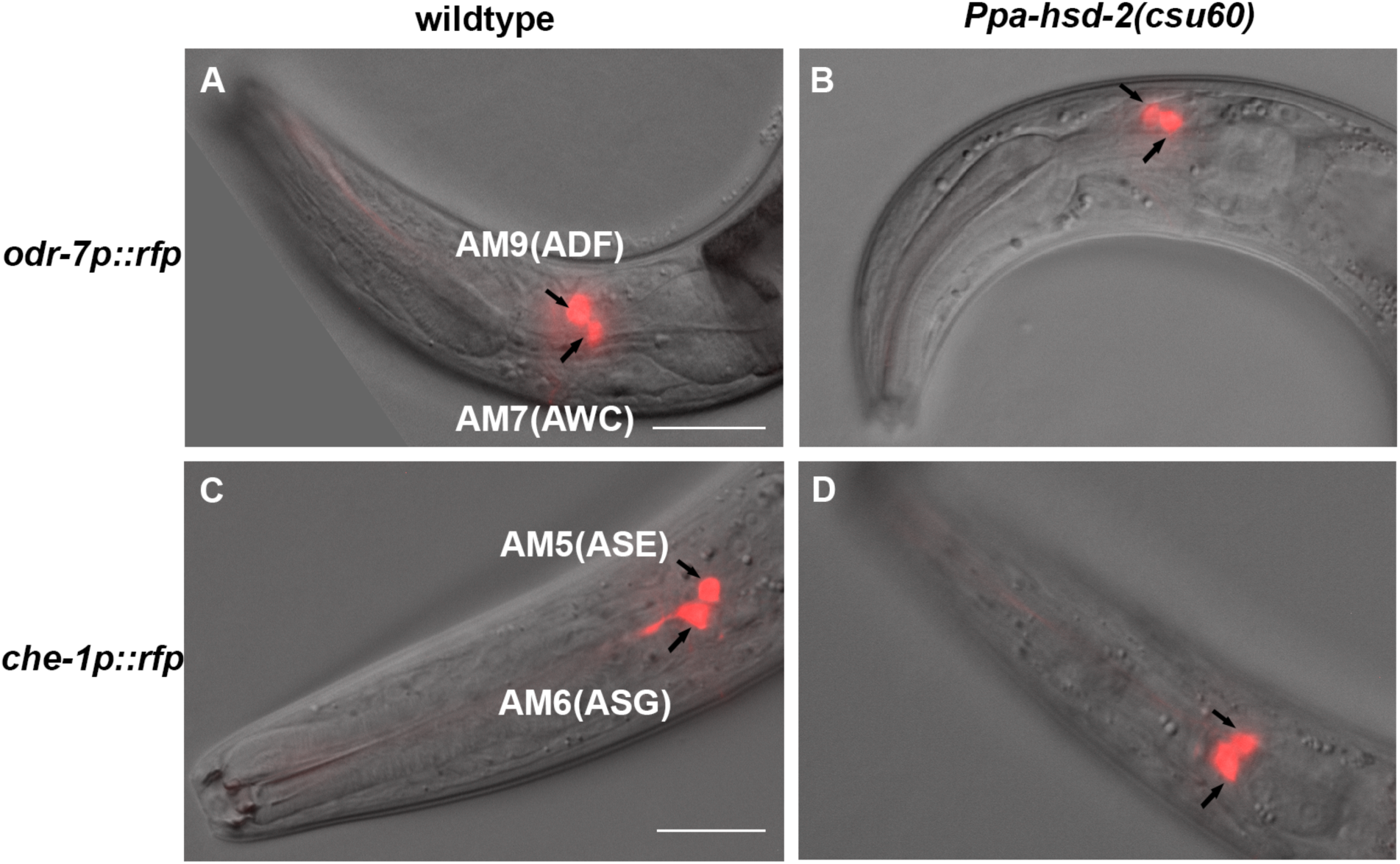
**Amphid neuron marker expressions for two reporters are wildtype in *Ppa-hsd-2(csu60)***. Representative overlay images of adult hermaphrodites expressing the *Ppa-odr-7p::rfp* reporter in AM9(ADF) and AM7(AWC) neurons of **(A)** wild-type PS312 and **(B)** *hsd-2(csu60)* mutant. Overlay images of adult hermaphrodites expressing the *Ppa-che-1p::rfp* reporter in AM5(ASE) and AM6(ASG) neurons of **(C)** wild-type PS312 and **(D)** *hsd-2(csu60)* mutant. Anterior is left and dorsal is up. Scale bars represent each reporter strain: 20 µm.

**SI Figure 7.**
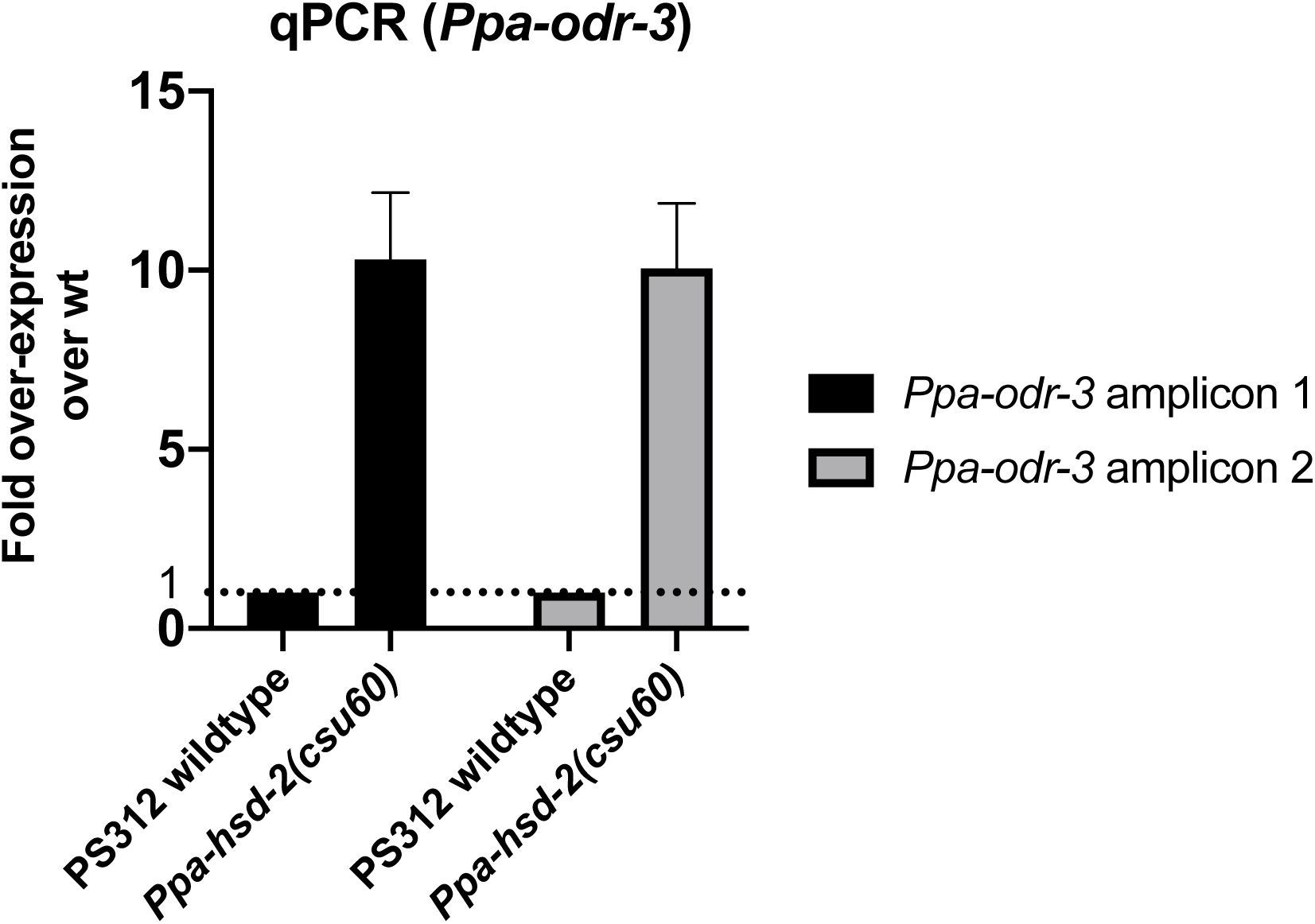
**RT-qPCR results of *Ppa-odr-3* expression**. Over-expression of *Ppa-odr-3* in *hsd-2(csu60)* is ∼10-fold more abundant than wild-type PS312 (arbitrarily set as one). Data represents 12 technical replicates from two biological RNA extraction replicates. Error bars denote s.e.m.

**SI Figure 8.**
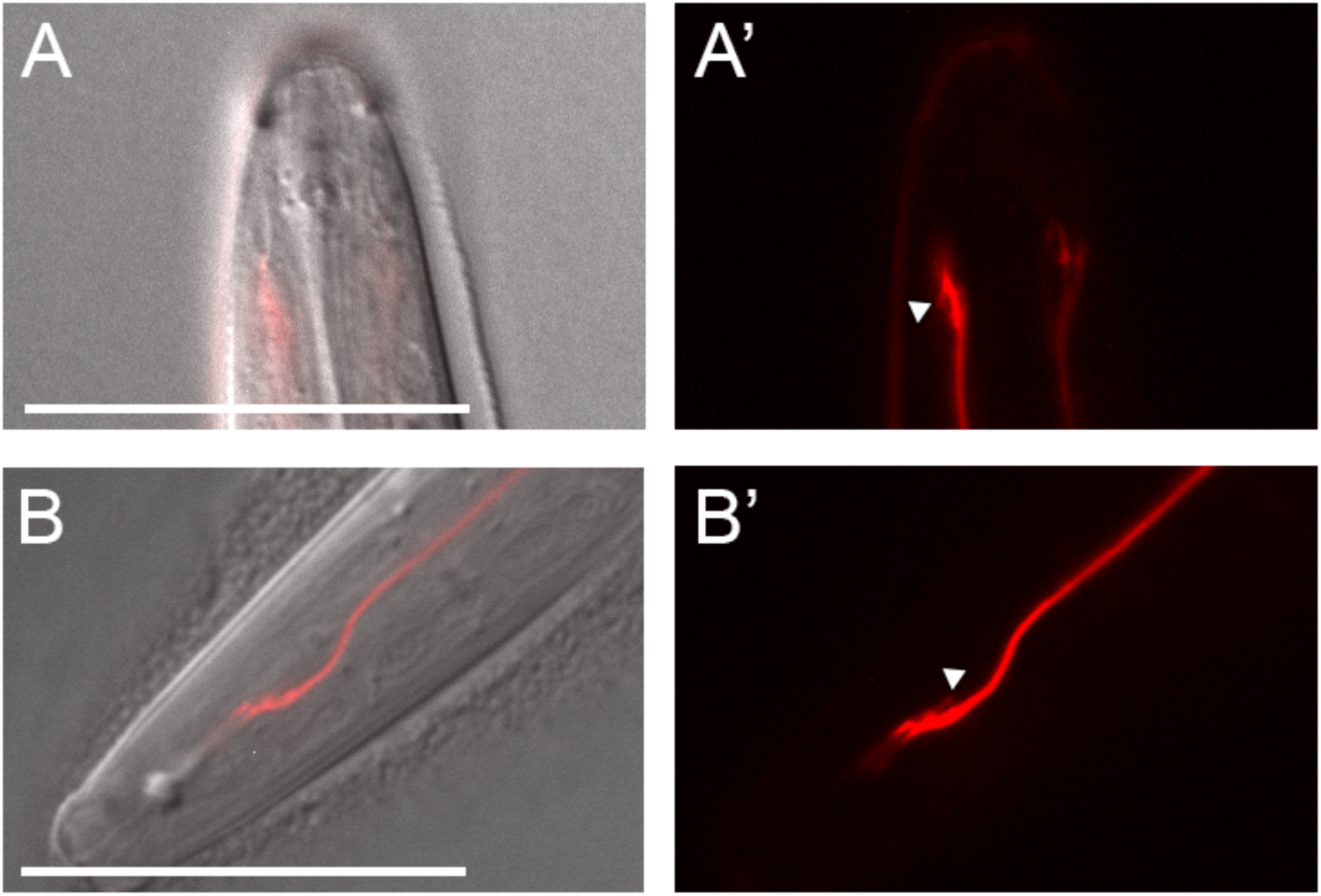
**Posterior protrusions in *Ppa-odr-3p::rfp* dendritic endings of *Ppa-hsd-2(csu60)* dauer larvae**. **(A-B)** Overlay images of additional examples of unusual dendritic ends found in *csu60* dauer larvae. **(A’ and B’)** RFP fluorescence. Triangle points to a type of posterior protrusion observed in 3 out of 17 *csu60* DL but not in wild-type DL. Anterior is up for (A), which is also a ventral view, and left for (B). Scale bar: 25 µm.

